# Brain-wide Activity Mapping Reveals the Somatosensory Cortex as a Sex-Specific Regulator of High-Fat Diet Intake

**DOI:** 10.64898/2026.03.24.714062

**Authors:** Chase A. Carter, Matthew T. Weaver, Samhitha S. Pudipeddi, Pierre Llorach, Jessica J. Walsh, Daniel J. Christoffel

## Abstract

High-fat diet (HFD) consumption engages reward circuitry and promotes neuroadaptations that contribute to overeating and obesity. While mesolimbic dopamine pathways are central to hedonic feeding models, the contribution of sensory cortical systems remains poorly understood. Here, we performed whole-brain activity mapping using Targeted Recombination in Active Populations (TRAP) and network analysis to define the distributed neural consequences of short-term HFD exposure in male and female mice.

HFD increased caloric intake in both sexes, with females consuming significantly more than males. Brain-wide analysis revealed striking sex-specific adaptations: HFD selectively increased isocortical activity in males, with the somatosensory cortex (SS) emerging as the most prominently modulated region. SS activity negatively correlated with HFD intake, primarily in males. Network analysis using the SMARTTR pipeline demonstrated that HFD reorganized network activity in a sex-dependent manner, biasing male networks toward associative cortical-thalamic hubs, whereas female networks preferentially recruited subcortical and brainstem structures.

To determine causality, we bidirectionally manipulated SS pyramidal neurons using chemogenetics during limited-access HFD exposure. Inhibition of the SS increased HFD intake in males, whereas activation reduced cumulative intake in females, without affecting locomotion. These findings establish the SS as a sex-specific regulator of palatable food consumption and demonstrate that similar behavioral outcomes emerge from distinct circuit architectures across sexes.

Collectively, this study expands prevailing reward-centric models of hedonic feeding by identifying sensory cortical control as a critical component of diet-induced neuroadaptations, with important implications for sex-specific therapeutic strategies targeting overeating and obesity.

## INTRODUCTION

The increasing prevalence of obesity and its adverse health consequences represents a major global public health concern. Excess adiposity is associated with cardiovascular disease, hypertension, hormone dysregulation, sleep disturbance, and numerous additional comorbidities.^1-3^ Since 1990, the prevalence of obesity among adults has approximately doubled worldwide,^4,5^ a trend that parallels increased consumption of energy-dense foods high in dietary fat.[6] This is highlighted by a recent meta-analysis that found consumption of ultra-processed foods, commonly enriched in sugars and fats, is associated with a 26% increased risk of obesity.[7] Importantly, high-fat diets (HFD) can induce neural adaptations similar to those produced by other highly rewarding stimuli.^8,9^ These diet-induced changes in the brain are thought to contribute to persistent overeating and the difficulty many individuals experience in maintaining healthy eating habits observed in obesity and related eating disorders.^10^

Food consumption is commonly conceptualized through two partially overlapping regulatory frameworks: homeostatic feeding and hedonic feeding. Homeostatic feeding supports metabolic needs and is primarily regulated by hypothalamic and brainstem circuits where neurotransmitters and hormonal signals mediate communication between the brain and gastrointestinal tract.^11^ In contrast, hedonic feeding reflects the rewarding and motivational properties of food and often occurs independent of metabolic necessity. This form of feeding is widely attributed to mesolimbic dopamine circuitry, particularly projections from the ventral tegmental area to the nucleus accumbens, amygdala, prefrontal cortex, and hippocampus.^12-14^ Within this framework, palatable foods engage reward systems in ways that resemble drugs of abuse, producing neuroadaptations that may reinforce compulsive intake.

Despite this emphasis on canonical reward circuitry, other brain regions involved in sensory processing may play an underappreciated role in feeding behavior. One candidate region is the somatosensory cortex (SS). While traditionally viewed as a relay node of sensory signals, recent evidence indicates that sensory cortices participate in adaptive decision-making and behavioral control.^15^ The SS processes tactile, thermal, and texture-related sensory features that are fundamental components of food perception and evaluation. Additionally, activity within oral somatosensory regions can influence orofacial movements required for ingestion, including movements of the mouth and tongue during consummatory behavior.^16^ Human neuroimaging studies further suggest that individuals prone to obesity exhibit altered somatosensory cortical responses during short-term energy imbalance, indicating that SS activity may be sensitive to metabolic state and food-related sensory cues.^17^

Given that feeding behavior emerges from coordinated activity across distributed neural circuits, identifying brain-wide activity patterns associated with palatable food consumption is essential for understanding mechanisms that promote overeating. To address this, we performed an unbiased screen of neuronal activity at the time of HFD exposure and intake using Targeted Recombination in Active Populations (TRAP) to genetically label neurons engaged during HFD intake, followed by atlas-based whole-brain activity mapping. This approach identified the isocortex, particularly the SS, as a prominently modulated region during HFD exposure. Notably, these effects were sex-specific, with HFD primarily altering isocortical activity in male mice. Based on these observations, we hypothesized that SS activity contributes to the regulation of palatable food consumption. To test this, we performed chemogenetic manipulations and demonstrated SS activity causally influences HFD intake. Together, these studies identify the SS as a potential regulator of hedonic feeding and reveal sex-specific circuit adaptations associated with palatable diet exposure.

## MATERIALS AND METHODS

### Experimental design and blinding

All experiments were conducted in a blinded manner such that behavioral assays and analyses were performed without knowledge of experimental group. Animals were randomly assigned to experimental conditions using cage-based randomization.

### Animals

All transgenic mice were bred in-house on a C57Bl/6J background (*Fos^tm2*.*1(icre/ERT2)Luo^*/J, “TRAP”, Jackson Laboratory #030323; B6.Cg-*Gt(ROSA)26Sor^tm14(CAG-tdTomato)Hze^*/J, “Ai14”, Jackson Laboratory #007914). *TRAP2:Ai14* mice were maintained as double heterozygotes. Male and female mice were included in all experiments. Sex was treated as a biological variable and analyzed separately where indicated.

Mice were housed on a 12-h light/dark cycle (22°C and 40% humidity) with ad libitum access to food and water. Standard chow contains 21% protein, 64% carbohydrates, and 15% fat (4.30 kcal/g; PicoLab 5V5R). Body weight was measured prior to and following the conclusion of each experiment at a consistent time. No statistical methods were used to predetermine sample size. All procedures were approved by the University of North Carolina Institutional Animal Care and Use Committee and were conducted in accordance with NIH Guidelines for the Care and Use of Laboratory Animals.

### High-fat diet intake

Mice were group-housed during the HFD exposure period. HFD procedures were adapted from previous studies.^18,19^ Weight-matched mice were randomly assigned to experimental groups. For each exposure, mice were separated and acclimated to individual cages for 30 min prior to diet presentation. The HFD used to model binge-like intake consisted of 20% protein, 20% carbohydrates, and 60% fat (5.24 kcal/g; Research Diets D12492).

For TRAP experiments, days 1–3 consisted of saline (intraperitoneal, i.p.) injections followed by 30 min habituation and 1-h access to either a single, pre-weighed standard chow pellet from their home cage or a novel high-fat pellet at a consistent time each day. On day 4, mice received 4-hydroxytamoxifen (4-OHT; Sigma, #H6278; 10 mg/mL in 1:4 castor oil:sunflower seed oil, 0.01 mL/g body weight), 30 min prior to a 3-h diet exposure. Food intake was calculated by subtracting post-exposure pellet weight from pre-exposure weight, converting to calories, and normalizing to body weight (cal/g BW). For chemogenetic experiments, mice received an i.p. injection followed by 30 min habituation prior to 20 min access to a pre-weighed HFD pellet. Mice received an i.p. injection of saline on day 1 and deschloroclozapine (DCZ) on days 2–4.

### Histological preparation, whole-brain imaging, and cell quantification

Two weeks following 4-OHT administration, mice were perfused and brains were post-fixed in 4% paraformaldehyde. Tissue was sectioned using a vibratome (Leica 1000S), mounted and coverslipped with Fluoromount-G containing DAPI (Invitrogen). Coronal sections were imaged using a Keyence BZ-X800 microscope at 10× magnification. Images were registered to the Allen Brain Atlas Common Coordinate Framework (CCF) using ImageJ and Aligning Big Brains and Atlases (ABBA).^20^ tdTomato^+^ cells were automatically detected and quantified within atlas-defined regions using custom Fiji macros.

### Network analysis using SMARTTR

Whole-brain network analysis was performed using the Simple Multi-Ensemble Atlas Registration and Statistical Testing in R (SMARTTR) workflow pipeline.^21^ This approach applies graph theory-based methods to quantify functional relationships and topological properties across brain regions derived from whole-brain activity mapping. Region-wise activity values (tdTomato^+^ cell counts) were first extracted from atlas-defined regions following registration to the Allen Brain Atlas Common Coordinate Framework. For each experimental group, pairwise Pearson correlation coefficients were calculated between all region pairs to generate region-to-region correlation matrices, representing functional coupling of activity across the brain. To focus on robust functional connections, correlation matrices were thresholded using a stringent criterion (Pearson’s r ≥ 0.9 or ≤ −0.9, α = 0.01), generating adjacency matrices that defined network edges. Separate networks were constructed for each experimental condition, and analyses were performed in a sex-stratified manner.

Global network metrics were averaged across all nodes within each condition, while regional (node-level) metrics were used to identify highly connected or influential brain regions. Topology metrics were computed for each network using SMARTTR, including: (1) degree centrality, representing the total number of connections per node; (2) clustering coefficient, reflecting local interconnectedness among neighboring nodes; (3) global efficiency, defined as the inverse of the average shortest path length across the network; and (4) betweenness centrality, quantifying the extent to which a node lies along shortest paths between other nodes and influences information flow.

To assess activation patterns that can signify differences in functional connectivity between conditions, permutation-based analyses were performed. Correlation differences were calculated by subtracting HFD values from corresponding SC values and compared against a permuted null distribution. Group labels were randomly shuffled to generate null distributions of correlation differences. Only connections exceeding both absolute magnitude (Δr ≥ 1) and significance (p < 0.01) thresholds were retained for downstream analysis. Network visualization and statistical comparisons were conducted within the SMARTTR framework in R, allowing for integrated analysis of both global and regional network properties.

### Stereotaxic surgery

Mice were anesthetized with ketamine (100 mg/kg) and xylazine (10 mg/kg), secured in a stereotaxic frame, and the skull was exposed. Viral infusions (0.3 μL) were delivered to the SS using a 33-gauge Hamilton syringe at 0.1 μL/min (coordinates: A/P +0.8 mm, M/L ±3.1 mm, D/V −2.25 mm, 4° angle). Needles were left in place for 5 min following infusion before withdrawal.

### Viruses

Adeno-associated viruses (AAV) used in this study were purchased from UNC Vector Core and Addgene and included: pAAV2-CaMKIIα-eYFP (UNC Vector Core), pAAV2-CaMKIIα-HA-hM4D(Gi)-IRES-mCitrine (Addgene #50467-AAV2), pAAV2-CaMKIIα-mCherry (Addgene #114469-AAV2), and pAAV2-CaMKIIα-hM3D(Gq)-mCherry (Addgene #50476-AAV2). AAV titers ranged from 4.4 × 10^12^ to 2.4 × 10^13^ gc/mL.

### Chemogenetic stimulation

Four weeks following surgery, mice expressing a Designer Receptor Exclusively Activated by Designer Drugs (DREADD) or control virus received DCZ (30 µg/kg, i.p., Tocris) or saline vehicle ∼30 min prior to HFD exposure.

### Statistical analysis

All statistical analyses were performed using GraphPad Prism (v10, GraphPad Software) and R (v4.2 or later) for network-based analyses. Data are presented as mean ± SEM unless otherwise stated. All datasets were assessed for normality using Shapiro-Wilk tests where appropriate. Variance was assumed to be similar between groups, and where this assumption was violated, appropriate corrections were applied. Statistical significance was set at α = 0.05, unless otherwise stated.

For behavioral and cell quantification data, group differences were assessed using two-way analysis of variance (ANOVA) with factors including diet (SC vs. HFD) and sex (male vs. female) or treatment (DREADD vs. control), as appropriate. When repeated measures were collected across days, repeated-measures ANOVA was used. Significant main effects or interactions were followed by Šídák’s multiple comparisons post hoc tests. For comparisons between two groups, unpaired Student’s *t*-tests were used. When appropriate, paired *t*-tests were used for within-subject comparisons. Correlation analyses were performed using Pearson’s correlation coefficients (*r*) to assess relationships between neuronal activity and behavioral measures. Linear regression was used to evaluate goodness of fit where indicated. Differences in topology network metrics between groups were assessed using unpaired *t*-tests.

## RESULTS

### Mapping diet-induced activity patterns reveals sex-specific differences in response to HFD exposure

To determine how short-term limited-access HFD exposure influences brain-wide neuronal activity compared to standard chow (SC), we utilized *TRAP2^+/-^:Ai14^+/-^* transgenic mice to label endogenously active neurons. In this system, iCreERT2 expression is driven by the *Fos* promoter and is activated by 4-hydroxytamoxifen (4-OHT), enabling permanent recombination and expression of the fluorescent reporter tdTomato in neurons active during the labeling window.

To capture neuronal populations relevant to hedonic feeding, mice were exposed to either HFD or SC over four consecutive days. On day 4, mice received 4-OHT (50 mg/kg i.p.) 30 min prior to a 3-h diet exposure (Figure 1A). The extended exposure period was used to maximize labeling of neurons engaged during diet consumption. Following tissue collection, our activity mapping pipeline consisted of slide-scanner imaging of coronal brain sections followed by semi-automatic registration to the Allen Brain Atlas CCF using Aligning Big Brains and Atlases.^22^ Automated detection and quantification of tdTomato^+^ cells were then performed across atlas-defined regions of interest (Figure 1B).

**Figure 1.**
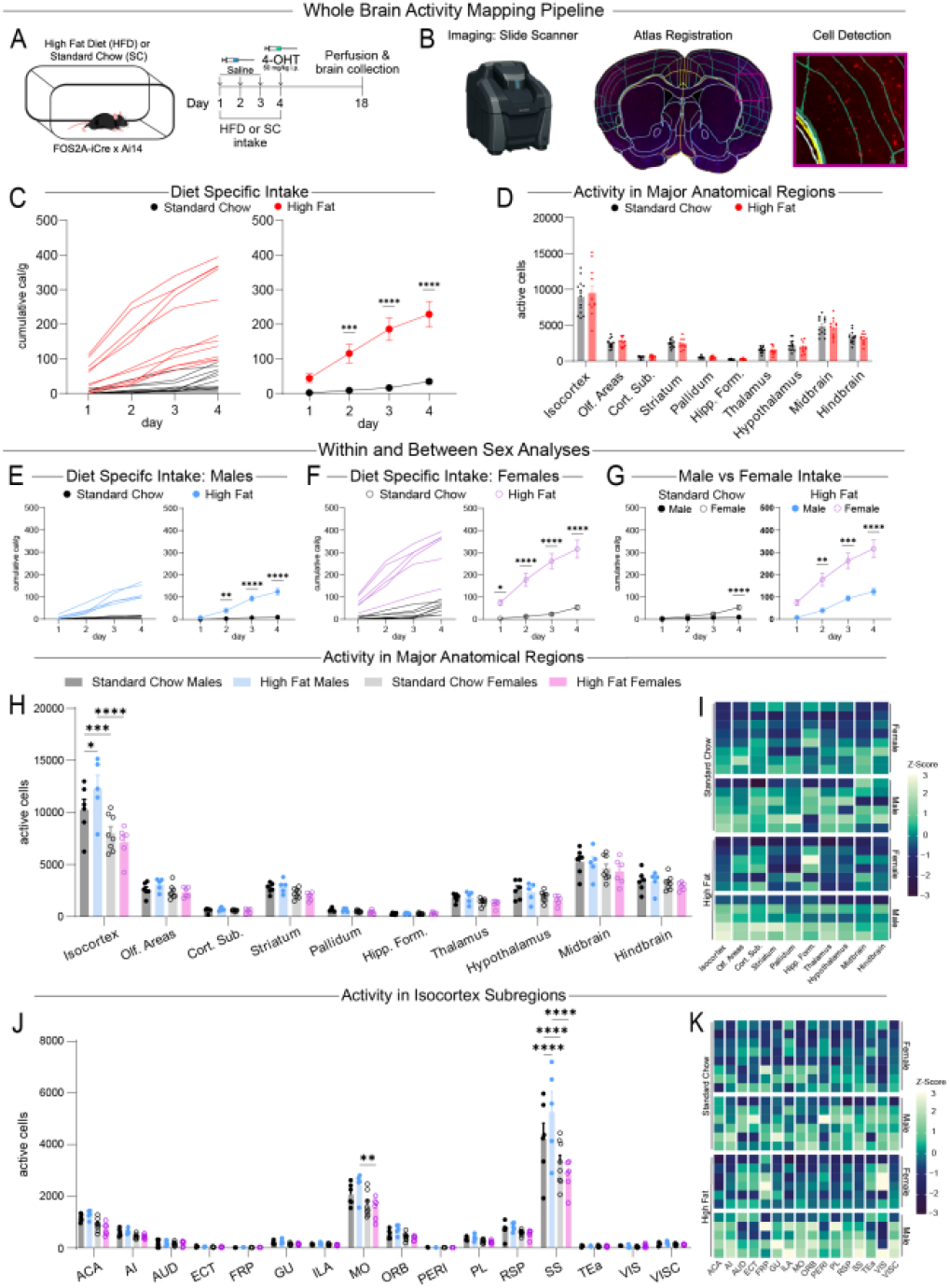
A Schematic of targeted recombination in active populations experiment for standard chow or high fat diet exposure (left) and experimental timeline (right). B Schematic of activity mapping pipeline. C Individual and average cumulative intake in cal/g (F(3, 69) = 38.24, p < 0.0001). D Active cell quantification of the major anatomical regions. E Male (F(3, 27) = 48.21, p < 0.0001) and F female (F(3, 36) = 43.75, p < 0.0001) individual cumulative intake in cal/g and averages. G Average sex cumulative intake in cal/g by standard chow (F(3, 36) = 10.73, p < 0.0001) and high fat diet (F(3, 27) = 10.24, p < 0.0001). H Active cell quantification of major anatomical regions by sex and diet (F(27, 210) = 3.219, p < 0.0001). I Heatmaps of individual mice by major anatomical regions. J Active cell quantification of the isocortex subregions by sex and diet (F(45, 336) = 3.831, p < 0.0001). K Heatmaps of individual mice by isocortex subregions. Data are shown as mean ± s.e.m. *p < 0.05, **p < 0.01**, *p < 0.001, ****p < 0.0001, two-way ANOVA with Šídák’s multiple comparison post hoc test. See Table 1 for full list of acronyms.

As expected across the 4 days of diet exposure, mice exposed to HFD consumed significantly more calories per gram body weight than SC controls (Figure 1C). When sexes were combined, however, we found no significant differences in the number of active cells across major anatomical regions between HFD and SC groups (Figure 1D). Sex-stratified analyses confirmed that both males and females consumed more HFD than SC (Figure 1E–F), and that females consumed approximately threefold more HFD than males across the 4-day exposure period (Figure 1G).

Given this pronounced behavioral difference, we next performed sex-specific quantification of activity patterns. HFD exposure selectively altered activity within the isocortex in male but not female mice. In addition, baseline sex differences in isocortical activity were observed between male and female mice exposed to SC. No significant differences in active cell counts were detected across major anatomical regions in female mice exposed to HFD compared with SC (Figure 1H–I). To further resolve regional differences, we next analyzed activity at the level of Allen CCF subregions (Figure 1J–K; Figure S2, S3; Table 1). Within the isocortex, the somatosensory cortex (SS) emerged as the most prominently modulated region. Male mice exposed to HFD exhibited increased SS activity compared with SC males, and consistently higher activity in males compared to females across conditions (Figure 1J–K).

**Table 1.**
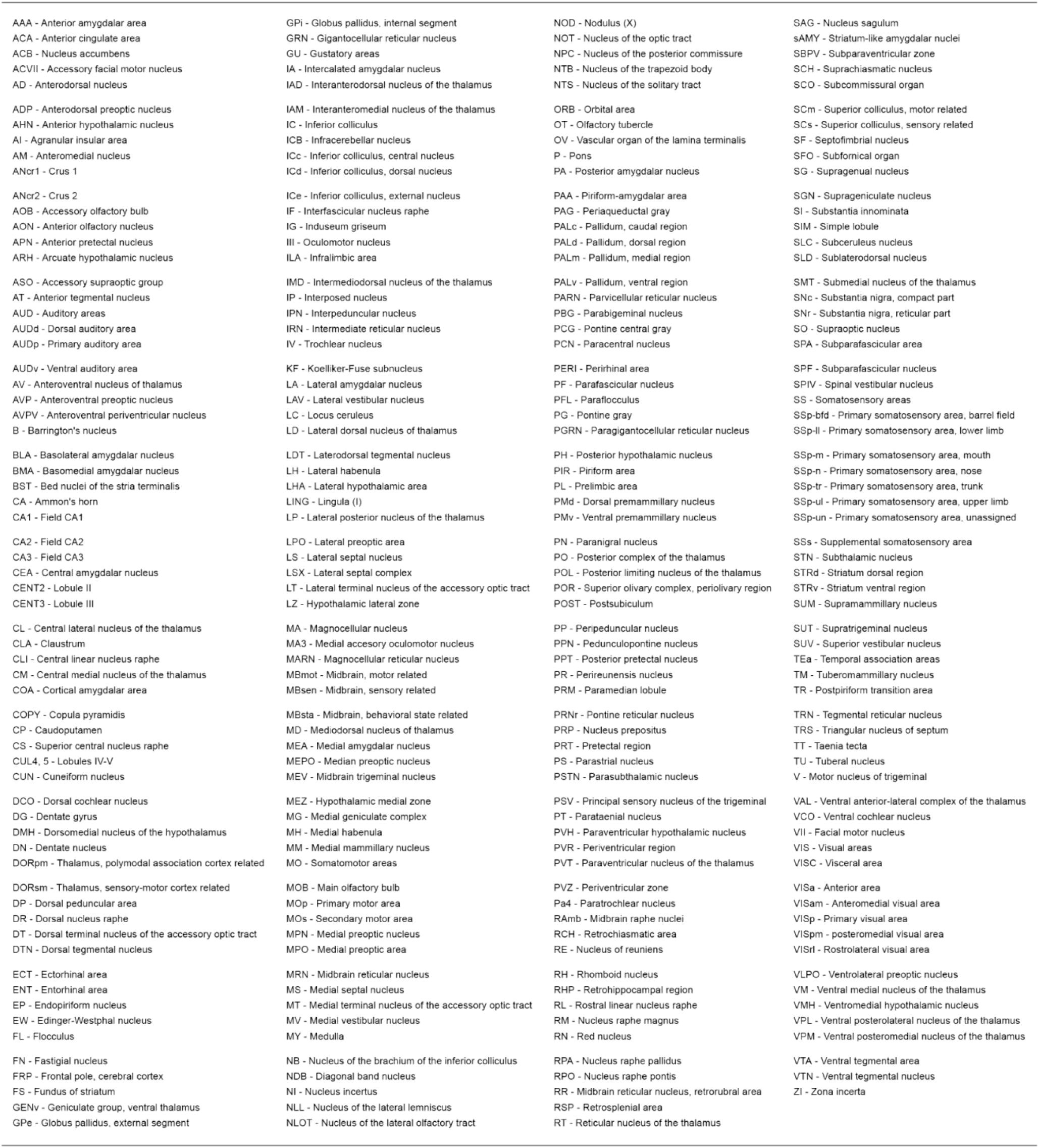
Allen Brain Atlas Common Coordinate Framework regions of interest list with acronym and full name.

During the TRAP labeling window, HFD intake remained elevated compared to SC intake (Figure S1A–C), and SS activity was negatively correlated with caloric consumption, an effect primarily driven by male HFD mice (Figure S1D–F). Additional region-specific differences were observed in olfactory and amygdala subregions (Figure S2), though no other major anatomical regions showed consistent diet-dependent effects (Figure S3). These findings implicate the isocortex, specifically the SS, in sex-specific modulation of hedonic feeding.

### Network analysis of diet-induced activity patterns reveals sex-specific differences in response to HFD exposure

To determine how HFD alters large-scale brain activity patterns, we performed network analysis using the SMARTTR workflow pipeline^21^ in a sex-stratified manner. SMARTTR enables visualization and quantification of network topology using metrics including degree centrality (overall connectivity of a brain region or “node”), clustering coefficient (interconnectedness among nodes sharing a common neighbor), efficiency (inverse of the average shortest path length), and betweenness centrality (influence over information flow within the network). We first constructed region-to-region correlations representing strong positive or negative coupling in activity as an index of functional connectivity between brain regions.

In male mice, HFD exposure increased positive correlations and reduced negative correlations across all brain regions (Figure 2A, B; average correlation: SC = 0.043, HFD = 0.234), and after applying a stringent threshold to identify robust region-to-region associations (Figure 2C–D; positive correlations: SC = 396, HFD = 508; negative correlations: SC = 156, HFD = 80). Global network topology analysis revealed increased clustering coefficient and reduced efficiency in HFD networks, with no change in degree or betweenness centrality (Figure 2E–I), suggesting enhanced local connectivity alongside reduced global integration.

**Figure 2.**
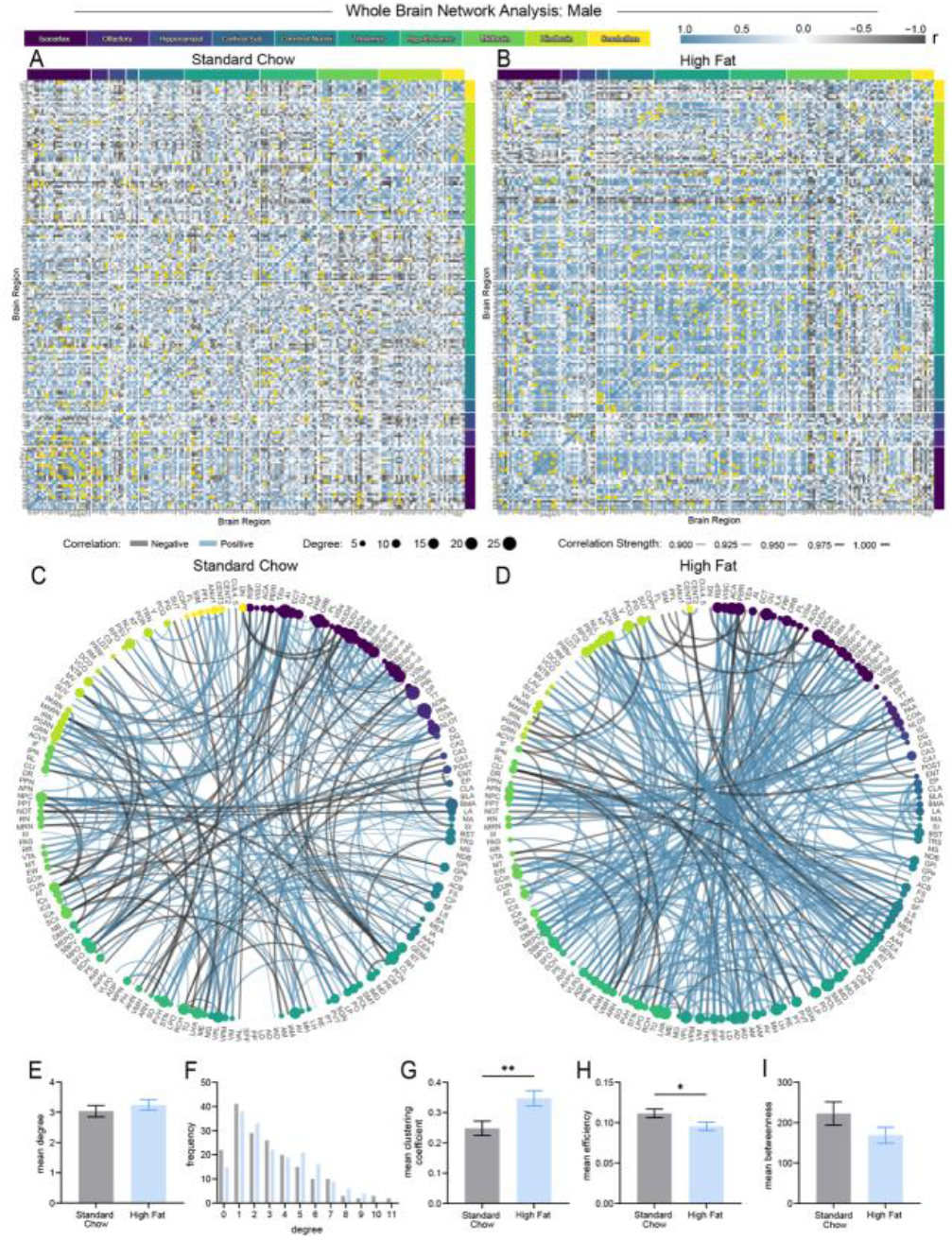
Male whole-brain correlational analysis of standard chow and high fat diet by utilizing SMARTTR. A Standard chow and B high fat diet Allen Brain Atlas subregion heatmaps of negative (gray; SC = 0.471, HFD = 0.314) and positive (blue; SC = 0.529, HFD = 0.686) correlations. C Standard chow and D high fat diet network analyses showing negative (gray; SC = 156, HFD = 80) and positive (blue; SC = 396, HFD = 508) between subregion correlations (display criteria: Pearson-r > 0.9 or < −0.9 and α = 0.01). Network attributes by standard chow (gray) and high fat diet (blue): E–F degree mean (t = 0.8198, df = 364, p = 0.4128) and frequency, G mean clustering coefficient (t = 2.864, df = 364, p = 0.0044), H mean efficiency (t = 2.153, df = 364, p = 0.0320), and I mean betweenness (t = 1.541, df = 364, p = 0.1243). Data are shown as mean ± s.e.m. *p < 0.05, **p < 0.01, unpaired Student’s t-test.

To determine which specific brain regions contributed to these differences, we examined regional topology metrics (Figure S4A–D). In SC networks, node degree analysis highlighted regions associated with sensory valuation, context-dependent learning, and action planning (Figure S4A). The top ten regions were largely cortical and included areas involved in context-guided actions, decision-making,^23-25^ and modality-specific sensory processing.^26-30^ In contrast, HFD networks enriched for associative cortical and thalamic hubs (Figure S4A–D). Regional degree analysis identified regions associated with context-dependent action selection (MOs), associative and emotional memory (PERI), taste processing (GU), and somatosensory integration (VPL), suggesting engagement of circuits that integrate sensory valuation with contextual and action-related signals during HFD exposure.^31-35^ Further, hypothalamic regions, including the lateral hypothalamic area (LHA) and tuberal nucleus (TU), were present, consistent with their established roles in feeding regulation and reward seeking.^36-39^ (Figure S4B).

Complementary analysis of betweenness centrality in SC networks identified several regions acting as hubs for information routing (Figure S4D). These regions are associated with internal state regulation, oculomotor control, memory, as well as arousal and attentional processes.^40-43^ Together, these findings suggest that SC networks emphasize sensory and contextual systems supporting adaptive orienting behaviors. The HFD network exhibited a partially overlapping but functionally distinct set of influential hubs (Figure S4D). PERI emerged as the strongest node, followed by ventromedial thalamus (VM), orbitofrontal cortex (ORB), retrosplenial cortex (RSP), and primary motor cortex (MOp), areas implicated in spatial and emotional memory, decision-making, and reward.^44-47^ To further examine regional connectivity patterns, we analyzed correlation distributions for the top four regions in the HFD condition. This revealed a shift toward negative correlations in PERI, whereas the VM region showed a shift toward positive correlations under HFD conditions. Correlation distributions for the ORB and RSP were similar across diet conditions (Figure S4E). Altogether, these regions are well positioned to coordinate information flow between sensory, limbic, and motor systems, suggesting that HFD network states involve both widespread connectivity and selective routing through associative cortical and basal ganglia-related hubs.^31,48,49^

Finally, we performed permutation analysis to directly assess differences in network connectivity between HFD and SC conditions. This analysis revealed a greater number of altered connections occurring in the positive direction in HFD compared with SC networks (Figure S4F). Notably, several correlations that were negative under SC conditions became positive in HFD networks. These included connections among regions involved in feeding regulation, homeostatic signaling, and reward processing, including the prelimbic cortex (PL), hippocampal CA1, lateral habenula (LH), subfornical organ (SFO), and median preoptic nucleus (MEPO).^31,33,41,50-53^ Additional altered connections involved thalamic, sensory, and motor-related regions associated with the perception and consumption of HFD (Figure S4G).

In contrast to males, female mice exhibited a distinct pattern of network reorganization. HFD exposure increased the proportion of negative correlations without altering global topology metrics (Figure 3A–I). Specifically, across all brain regions HFD-exposed animals displayed a greater proportion of negative correlations compared with SC controls (average correlation: SC = 0.175, HFD = 0.124; Figure 3A–B). After applying a stringent threshold to identify robust region-to-region associations, HFD females exhibited 132 negative correlations, compared with 68 in SC females (Figure 3C–D).

**Figure 3.**
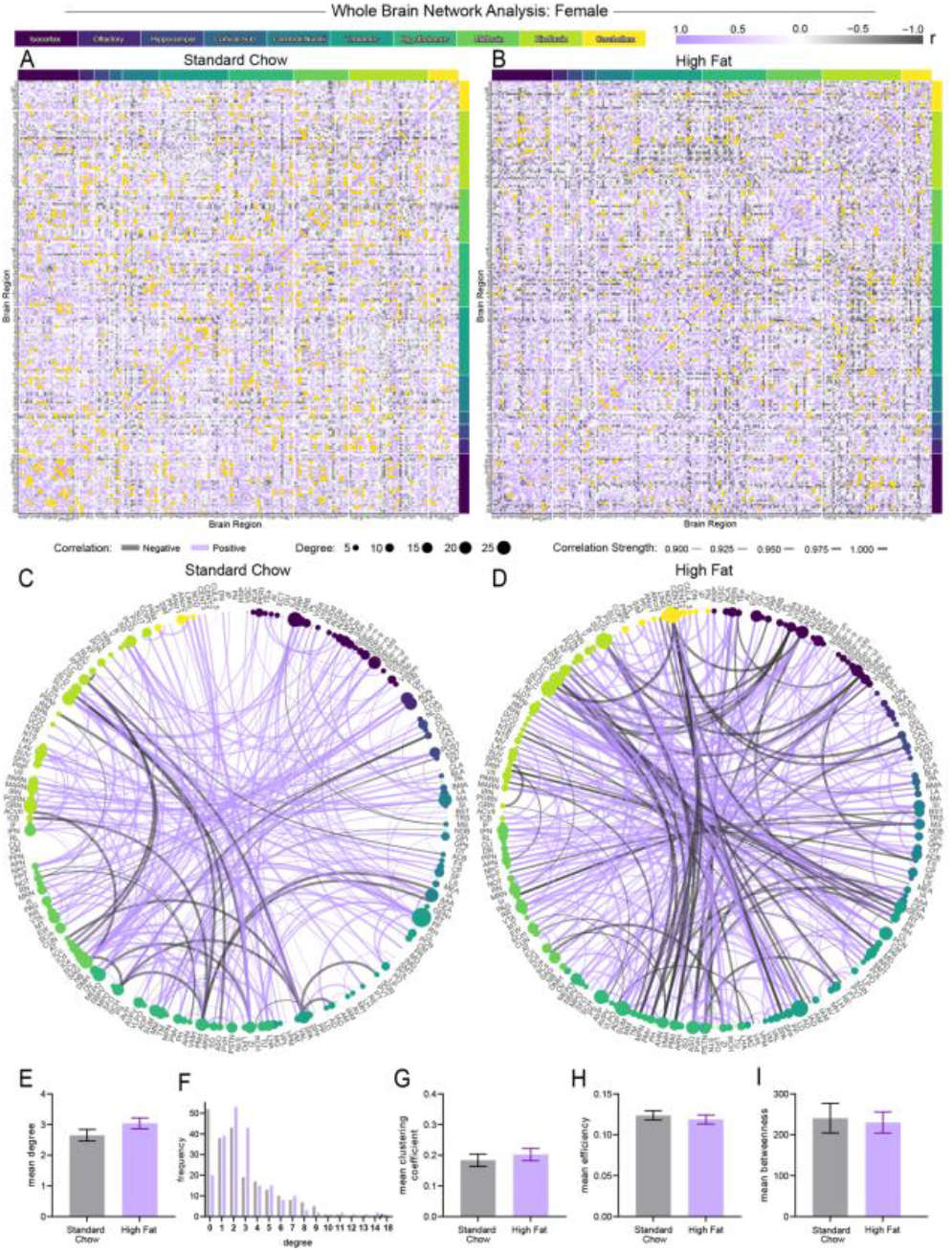
Female whole-brain correlational analysis of standard chow and high fat diet by utilizing SMARTTR. A Standard chow and B high fat diet Allen Brain Atlas subregion heatmaps of negative (gray; SC = 33.8%, HFD = 39.1%) and positive (purple; SC = 66.2%, HFD = 60.9%) correlations. C Standard chow and D high fat diet network analyses showing negative (gray; SC = 68, HFD = 132) and positive (purple; SC = 502, HFD = 520) between subregion correlations (display criteria: Pearson-r > 0.9 or < −0.9 and α = 0.01). Network attributes by standard chow (gray) and high fat diet (purple): E–F degree mean (t = 1.515, df = 428, p = 0.1305) and frequency, G mean clustering coefficient (t = 0.636, df = 428, p = 0.5251), H mean efficiency (t = 0.632, df = 428, p = 0.5279), and I mean betweenness (t = 0.237, df = 428, p = 0.8132). Data are shown as mean ± s.e.m. Unpaired Student’s t-test.

Regional analyses revealed that HFD networks in females were dominated by subcortical, brainstem, and cerebellar structures, including hypothalamic, pontine, and locus coeruleus nodes (Figure S5A–D). In SC females, high degree nodes included regions within the midbrain and olfactory structures, which respond to multimodal sensory and olfactory environmental cues,^54-56^ as well as frontal cortex and amygdala areas associated with context-dependent emotional processing.^57-59^ Additional regions included hindbrain areas involved in locomotor and head movement control,^42,60^ and hypothalamic, thalamic, and cortical areas associated with fluid consumption, attentional processes, and wakefulness.^40,41,61-64^

Female HFD network analysis identified thalamic, cerebellar, and pontine nuclei, along with multiple hypothalamic regions as highly connected nodes. The locus coeruleus (LC) also emerged as a prominent hub, consistent with its role in arousal, attention, and affective modulation (Figure S5A).^68^ Clustering analysis further identified highly interconnected regions associated with sensory processing, decision-making, and feeding and reward-related behaviors,^34,69-74^ while network efficiency remained similar between diet conditions (Figure S5B–C).

Analysis of influential information flow (betweenness centrality) nodes revealed similar functional characteristics (Figure S5D). Top regions included thalamic areas associated with attention, arousal, and auditory processing,^62,75^ along with hypothalamic regions that regulate homeostatic and goal-driven feeding and thirst signals.^41,76,77^ These analyses indicate that SC female networks emphasize systems involved in sensory valuation, contextual decision-making, and homeostatic regulation.

In comparison, HFD female networks again showed stronger influence of subcortical control structures as information-routing hubs. High betweenness nodes included cerebellar, pontine, medullary, and midbrain structures. Hypothalamic nuclei associated with feeding behavior,^77-80^ including the parasubthalamic nucleus (PSTN) and dorsal premammillary nucleus (PMd), were also prominent (Figure S5D). These patterns suggest that female HFD networks preferentially engage subcortical circuits involved in autonomic regulation and motor readiness, with comparatively reduced involvement of associative cortical hubs. Among the four regions with the highest betweenness values, correlation distributions revealed a shift toward negative correlations in the cerebellar nodulus (NOD), whereas the supragenual nucleus (SG) showed a modest shift toward positive correlations in HFD conditions (Figure S5E).

Permutation analysis indicated that HFD preferentially strengthened negative correlations among circuits involved in homeostatic regulation, valuation, and action selection (Figure S5F). These changes included altered correlations among regions involved in homeostatic regulation, reward valuation, contextual memory, and action selection,^45,51,81-86^ including the paraventricular hypothalamic nucleus (PVH), PSTN, ORB, hippocampal CA1, nucleus accumbens (ACB), and caudate putamen (CP) (Figure S5G). Together, these results demonstrate that HFD produces sex-specific reorganization of brain-wide networks, with males engaging associative cortical-thalamic systems and females preferentially recruiting subcortical and autonomic circuits.

### Somatosensory cortex regulation of HFD consumption

Given the sex-specific modulation of isocortical and SS activity, as well as the inverse relationship between SS activity and HFD consumption, we next examined the functional role of SS pyramidal neurons (SS^Pyr^) in regulating HFD intake. To do so, we utilized a chemogenetic approach to bidirectionally modulate SS^Pyr^ activity during short-term, limited-access HFD exposure. We bilaterally expressed either an inhibitory DREADD (AAV-CaMKIIα-hM4D(Gi)) or control virus (AAV-CaMKIIα-eYFP) selectively in excitatory pyramidal neurons of the SS (Figure 4A). On day 1, mice received a saline injection 30 min prior to HFD exposure to establish baseline intake. On days 2–4, mice received DCZ (30 µg/kg) prior to diet exposure (Figure 4B). When sexes were analyzed together, inhibition of SS^Pyr^ neurons through activation of G*i/o* signaling produced a modest but significant increase in HFD consumption, without affecting locomotor activity (Figure 4C–D). Based on our TRAP results, we next examined whether these effects differed by sex. Chemogenetic inhibition of SS^Pyr^ neurons produced a robust increase in daily HFD consumption in male mice, resulting in a significant increase in cumulative intake (Figure 4E), without altering locomotor behavior (Figure 4F). In contrast, inhibition of SS^Pyr^ neurons had no effect on HFD consumption or locomotion in females (Figure 4G–H).

**Figure 4.**
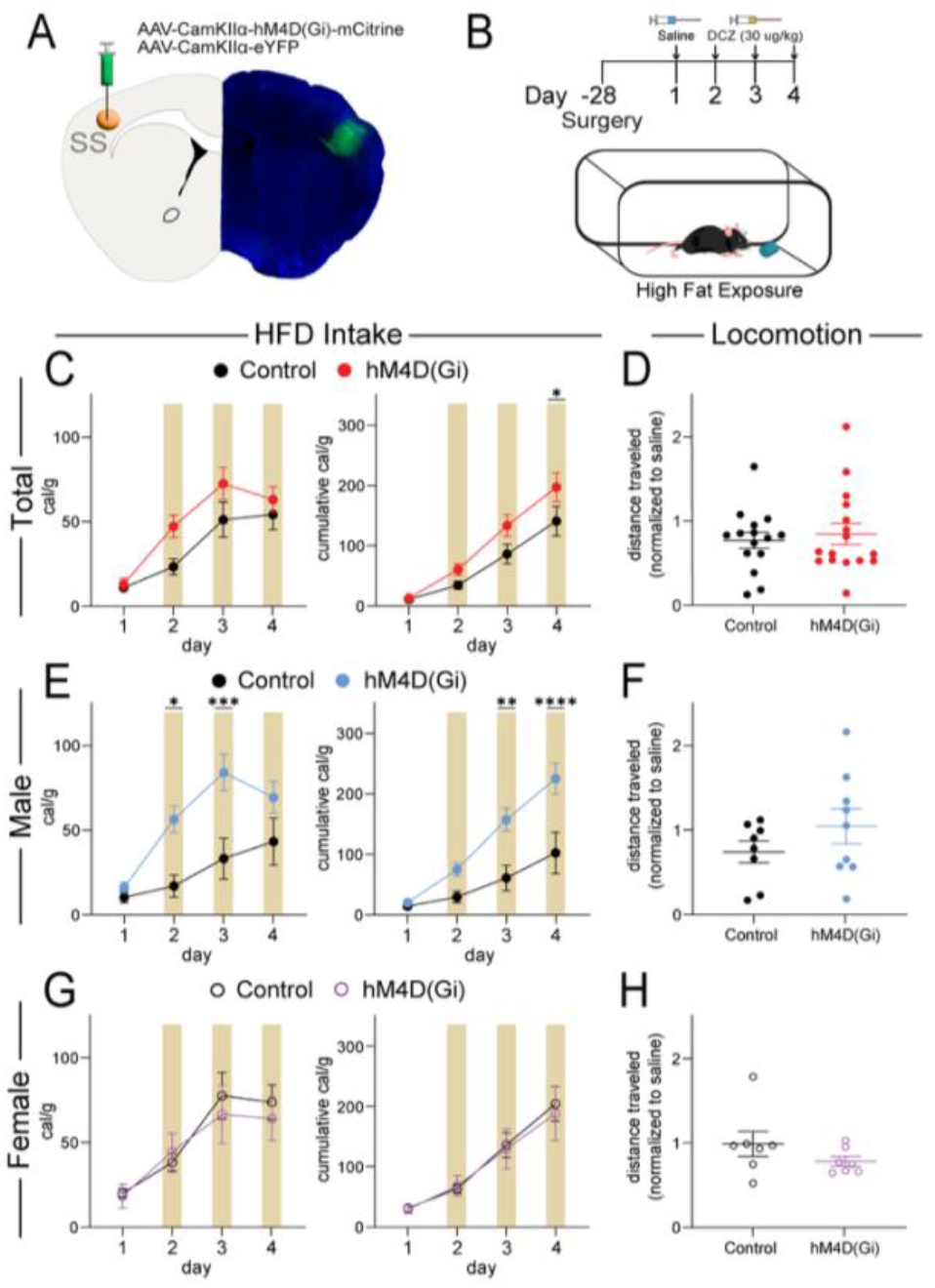
A Schematic of surgery for hM4D(Gi) DREADD expression (left) and representative image (right). B Schematic of high fat diet exposure experimental design. C Quantification of high fat diet intake with chemogenetic inhibition in all control (black) and hM4D(Gi) (red) expressing mice for daily (F(3, 90) = 2.453, p = 0.0684) and cumulative (F(3, 90) = 2.917, p = 0.0385) intake in cal/g. D Locomotion of all mice (t = 0.4846, df = 29, p = 0.6316). E Quantification of high fat diet intake with chemogenetic inhibition in male control (black) and hM4D(Gi) (blue) expressing mice for daily (F(3, 45) = 5.034, p = 0.0043) and cumulative (F(3, 45) = 8.773, p < 0.0001) intake in cal/g. F Locomotion of male mice (t = 1.210, df = 15, p = 0.2451). G Quantification of high fat diet intake with chemogenetic inhibition in female control (black) and hM4D(Gi) (purple) expressing mice for daily (F(3, 39) = 0.7000, p = 0.5577) and cumulative intake in cal/g (F(3, 39) = 0.1775, p = 0.9110). H Locomotion of female mice (t = 1.294, df = 12, p = 0.2201). Data are shown as mean ± s.e.m. *p < 0.05, **p < 0.01**, *p < 0.001, ****p < 0.0001, two-way ANOVA with Šídák’s multiple comparison post hoc test, and unpaired Student’s t-test.

In a parallel experiment using the same feeding paradigm (Figure 5A–B), we expressed an excitatory DREADD (AAV-CaMKIIα-hM3D(Gq)) in SS^Pyr^. Activation of SS^Pyr^ produced no significant changes in cumulative HFD intake or locomotion when sexes were combined, compared to control animals (AAV-CaMKIIα-mCherry) (Figure 5C–D). When analyzed separately by sex, male hM3D(Gq) mice showed no changes in consumption or locomotor activity relative to controls (Figure 5E–F). In females, acute activation of SS^Pyr^ on individual DCZ days did not significantly alter intake. However, cumulative HFD consumption was significantly reduced by the final day of HFD exposure (Figure 5G). Locomotor activity remained unchanged (Figure 5H). Together, these TRAP and chemogenetic experiments indicate that basal activity levels of the isocortex, particularly the SS, differ between the sexes and can influence palatable food consumption in a sex-specific manner.

**Figure 5.**
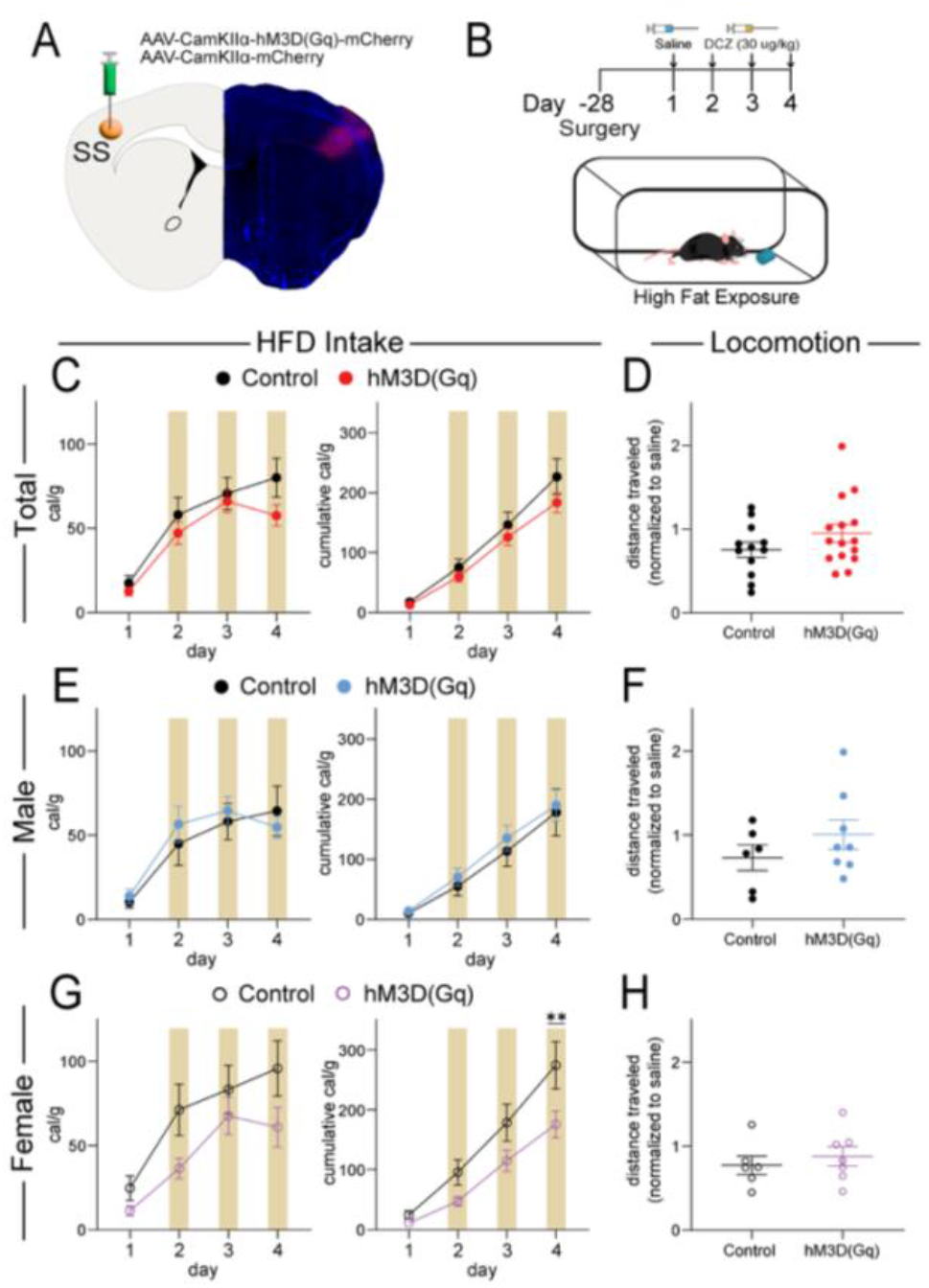
A Schematic of surgery for hM3D(Gq) DREADD expression (left) and representative image (right). B Schematic of high fat diet exposure experimental design. C Quantification of high fat diet intake with chemogenetic activation in all control (black) and hM3D(Gq) (red) expressing mice for daily (F(3, 75) = 1.195, p = 0.3176) and cumulative (F(3, 75) = 1.432, p = 0.2402) intake in cal/g. D Locomotion of all mice (t = 1.359, df = 25, p = 0.1863). E Quantification of high fat diet intake with chemogenetic activation in male control (black) and hM3D(Gq) (blue) expressing mice for daily (F(3, 36) = 1.239, p = 0.3099) and cumulative (F(3, 36) = 0.1779, p = 0.9107) intake in cal/g. F Locomotion of male mice (t = 1.137, df = 12, p = 0.2777). G Quantification of high fat diet intake with chemogenetic activation in female control (black) and hM3D(Gq) (purple) expressing mice for daily (F(3, 33) = 0.7756, p = 0.5160) and cumulative (F(3, 33) = 3.422, p = 0.0284) intake in cal/g. H Locomotion of female mice (t = 0.6484, df = 11, p = 0.5300). Data are shown as mean ± s.e.m. *p < 0.05, **p < 0.01**, *p < 0.001, ****p < 0.0001, two-way ANOVA with Šídák’s multiple comparison post hoc test, and unpaired Student’s t-test.

## DISCUSSION

In this study, we combined activity-dependent genetic labeling (TRAP), whole-brain mapping, network analysis, and chemogenetic manipulations to define the consequences of short-term HFD exposure on neural activity. Three principal findings emerged. First, HFD exposure produced robust, sex-specific alterations in isocortical activity, with the SS emerging as the most prominently modulated region in males. Second, HFD reorganized brain-wide functional networks in a sex-dependent manner, with males preferentially engaging associative cortical-thalamic hubs and females recruiting subcortical and brainstem structures. Third, causal manipulation of SS pyramidal neurons bidirectionally regulated HFD intake in a sex-specific manner. Together, these findings identify the SS as a previously underappreciated regulator of palatable food consumption and highlight sex as a critical determinant of diet-induced circuit adaptations.

The prevailing framework for hedonic feeding emphasizes mesolimbic dopamine circuitry, including projections from the ventral tegmental area (VTA) to the nucleus accumbens (ACB), ORB, amygdala, and hippocampus.^87-92^ Within this model, palatable food engages reward systems in a manner analogous to drugs of abuse, producing neuroadaptations that reinforce intake.^8,9,12,93-95^ Consistent with this literature, our network analyses identified canonical feeding and reward regions, including the orbitofrontal cortex (ORB), perirhinal cortex, thalamic nuclei, and hypothalamic structures such as the LHA, as central nodes in HFD-associated networks.^96-99^

A key finding of the present study is the selective recruitment of the isocortex, particularly the SS, in male mice exposed to HFD. SS activity was inversely correlated with caloric intake, suggesting that SS engagement may act to constrain consumption. This interpretation is supported by causal experiments: chemogenetic inhibition of SS pyramidal neurons increased HFD intake in males, whereas activation reduced cumulative intake in females. These findings extend the role of the SS beyond sensory relay, positioning it as an active participant in the regulation of feeding behavior.

The SS is well positioned to integrate tactile, thermal, and oral sensory signals with motor output during consumption.^29,100^ In humans, altered somatosensory responses have been reported in individuals prone to obesity during energy imbalance,^101^ suggesting that sensory cortical processing is sensitive to metabolic and motivational state.^110^ One parsimonious interpretation is that SS activity encodes sensory feedback that contributes to sensory-specific satiety or cortical gating of consummatory behavior. While cortical regions such as the ORB and insula are known to encode sensory valuation and termination of feeding,^12,96,102,103^ our findings identify the SS as an additional cortical node capable of exerting control over palatable intake.

Despite consuming substantially more HFD than males, female mice showed minimal isocortical modulation in response to diet. Instead, HFD networks in females were characterized by prominent engagement of subcortical, brainstem, cerebellar, and hypothalamic structures, including regions associated with arousal, autonomic regulation, and motor readiness. In contrast, male HFD networks were enriched for associative cortical and thalamic hubs. Permutation analyses further revealed sex-specific shifts in connectivity, with males showing increased positive coupling among regions involved in contextual processing and internal state regulation, and females showing enhanced negative coupling among homeostatic and action selection circuits.^14,96^ Furthermore, the piriform cortex and amygdala also exhibited diet- and sex-dependent modulation in our dataset and are known to integrate sensory cues with emotional and motivational valence.^104-106^ These findings indicate that similar behavioral outcomes can arise from distinct network architectures across sexes.

These sex-specific circuit organizations are consistent with broader evidence that sex influences reward processing, stress responsivity, and motivated behavior.^107-109^ However, most prior work has focused on subcortical reward systems, with relatively little attention to sensory cortical contributions. The present findings therefore extend current models of hedonic feeding by demonstrating that sensory cortical systems contribute to diet-induced circuit adaptations in a sex-dependent manner.

At the network level, using the SMARTTR workflow^21^ HFD exposure increased clustering while reducing efficiency, particularly in males, suggesting a shift toward greater local connectivity and more modular network organization. These changes may reflect coordinated recruitment of feeding-related ensembles while limiting global information transfer. Region-specific betweenness analyses further revealed that HFD networks rely on distinct hubs for information routing, with males favoring associative cortical nodes and females favoring subcortical structures. These results suggest that diet-induced neuroadaptations are not restricted to canonical reward pathways but instead involve distributed, sex-specific reorganization of large-scale brain networks.

Obesity and overeating are often conceptualized as disorders of dysregulated reward valuation or dopamine signaling.^89,110-113^ Our findings suggest that sensory cortical systems may exert top-down control over hedonic intake and that disruption of this control, particularly in males, can promote excessive consumption. This framework bridges homeostatic and hedonic models of feeding^91,97,114-116^ by positioning the SS at the interface of sensory feedback, motor output, and reward circuitry. Therapeutically, these data raise the possibility that targeting cortical sensory processing or circuit-level modulation may differentially influence feeding behavior across sexes.

### Limitations and future directions

Several limitations warrant consideration. First, TRAP labeling captures neuronal activity within a defined temporal window but does not resolve projection-defined or cell-type-specific subpopulations. Future studies using projection-specific approaches or *in vivo* recording will be necessary to determine how SS neurons interact with canonical feeding circuits, including the ACB and lateral hypothalamus. Second, the present study focuses on short-term HFD exposure, which models binge-like or acute palatable intake. Chronic diet exposure may produce distinct or progressive circuit adaptations relevant to long-term obesity and metabolic dysfunction. Third, while our findings establish a causal role for SS pyramidal neurons in regulating HFD intake, the downstream circuit mechanisms and synaptic substrates mediating these effects remain to be defined. Identifying the specific pathways through which SS influences feeding behavior will be an important direction for future work. Finally, although we identify robust sex differences in network organization and circuit function, the biological mechanisms underlying these differences, including hormonal, developmental, or genetic factors, remain unresolved. Elucidating these mechanisms will be critical for understanding how sex shapes vulnerability to maladaptive feeding.

## CONCLUSION

In summary, we identify the SS as a sex-specific regulator of HFD intake and demonstrate that palatable food exposure induces distinct, sex-dependent reorganization of brain-wide networks. These findings expand prevailing reward-centric models of hedonic feeding by incorporating sensory cortical control as a critical component of diet-induced neuroadaptation. More broadly, our results underscore the importance of integrating sex as a biological variable in studies of feeding behavior and suggest that similar behavioral outcomes may arise from fundamentally different circuit mechanisms across sexes.

## AUTHOR CONTRIBUTIONS

C.A.C., J.J.W., and D.J.C. designed the studies. C.A.C. and P.L. conducted the TRAP experiments. C.A.C., M.T.W., and S.S.P. conducted brain sectioning and imaging. C.A.C. and D.J.C. conducted cell counting, network analysis, and statistical analysis. C.A.C. and D.J.C. wrote the manuscript with feedback from M.T.W., S.S.P., P.L., and J.J.W.

## SUPPLEMENTAL MATERIALS

**Supplemental Figure 1.**
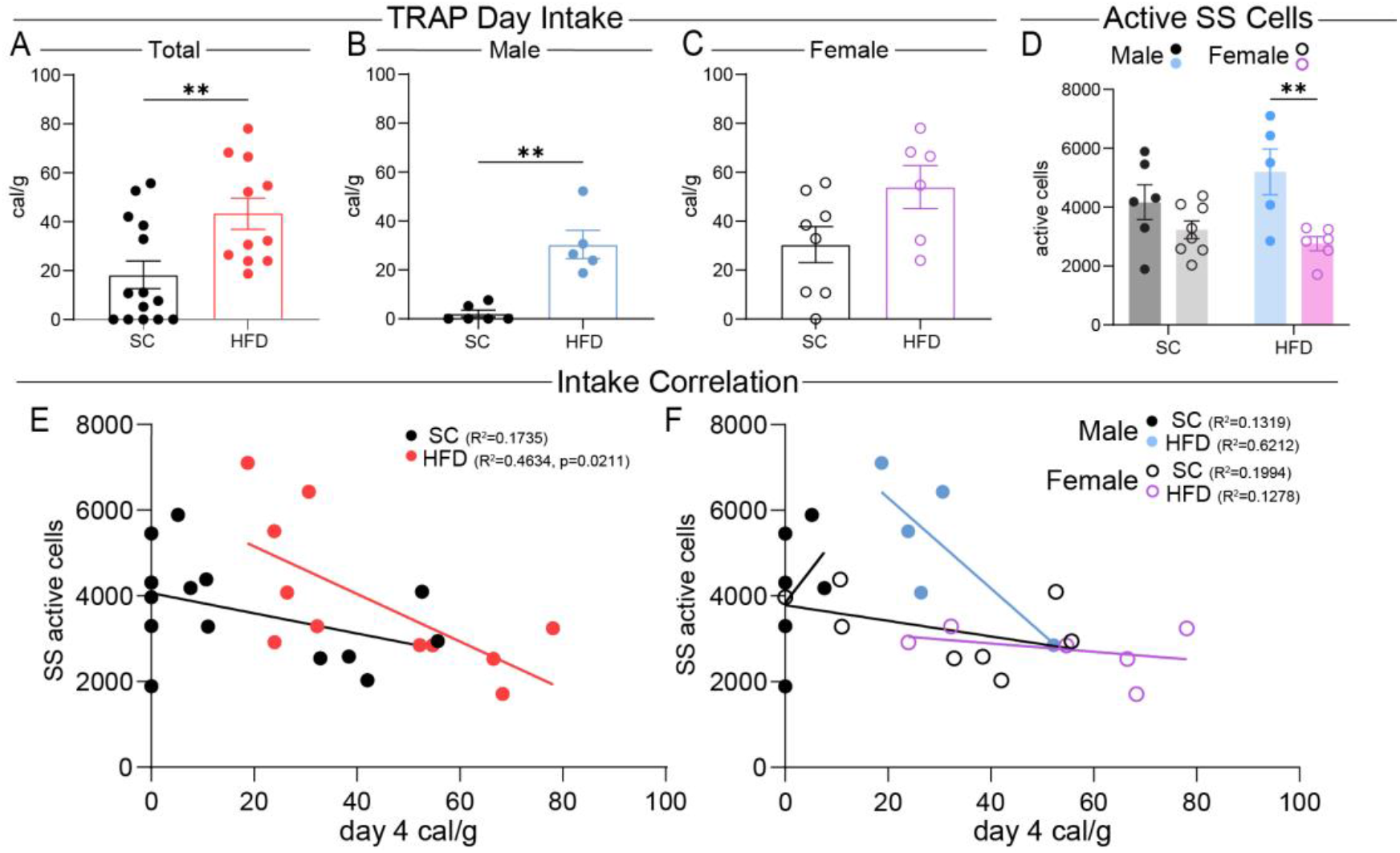
Targeted recombination in active populations consumption and activity correlations. Standard chow (black) and high fat diet (colored) day 4 consumption in cal/g for A total (t = 2.921, df = 23, p = 0.0077), B male (t = 5.186, df = 9, p = 0.0006), and C female (t = 2.066, df = 12, p = 0.0611). D Sex-delineated active somatosensory (SS) cells during day 4 consumption (F(3, 21) = 6.675, p = 0.0024). Intake correlation with active SS cells for E total (SC: R^2^ = 0.1735, p = 0.1384; HFD: R^2^ = 0.4634, p = 0.0211), F male (SC: R^2^ = 0.1319, p = 0.4791; HFD: R^2^ = 0.6212, p = 0.1133), and G female (SC: R^2^ = 0.1994, p = 0.2674; HFD: R^2^ = 0.1278, p = 0.4867). Data are shown as mean ± s.e.m. *p < 0.05, **p < 0.01**, *p < 0.001, ****p < 0.0001, one-way ANOVA, unpaired Student’s t-test, and linear regression.

**Supplemental Figure 2.**
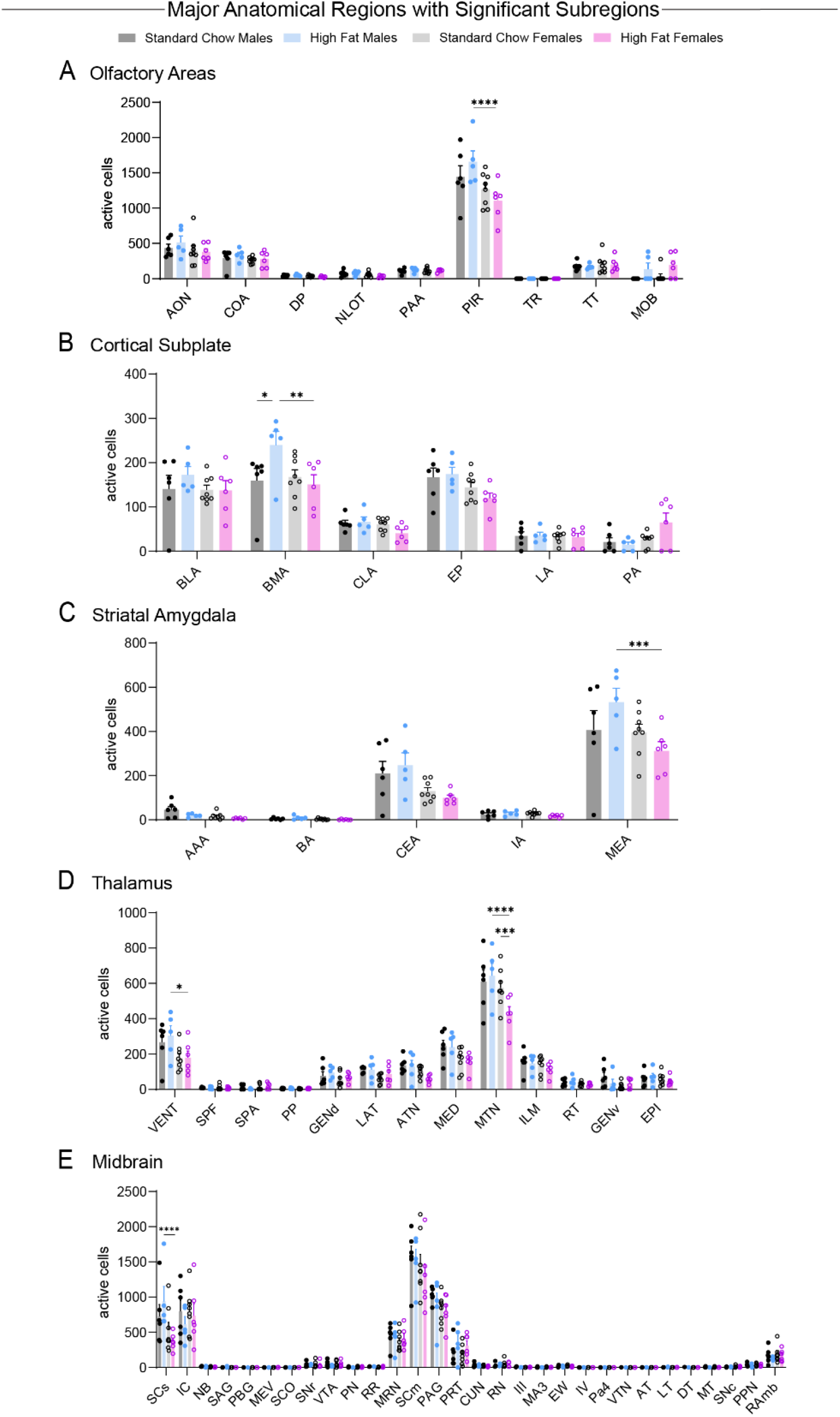
Active population cell quantification within major anatomical regions that contain significant subregions. A Olfactory Areas (F(24, 189) = 2.241, p = 0.0014). B Cortical Subplate (F(15, 126) = 1.736, p = 0.0519). C Striatal Amygdala (F(12, 105) = 1.723, p = 0.0720). D Thalamus (F(36, 273) = 1.698, p = 0.0102). E Midbrain (F(87, 630) = 1.069, p = 0.3238). Data are shown as mean ± s.e.m. *p < 0.05, **p < 0.01**, *p < 0.001, ****p < 0.0001, two-way ANOVA with Šídák’s multiple comparison post hoc test. Carter et al. 15

**Supplemental Figure 3.**
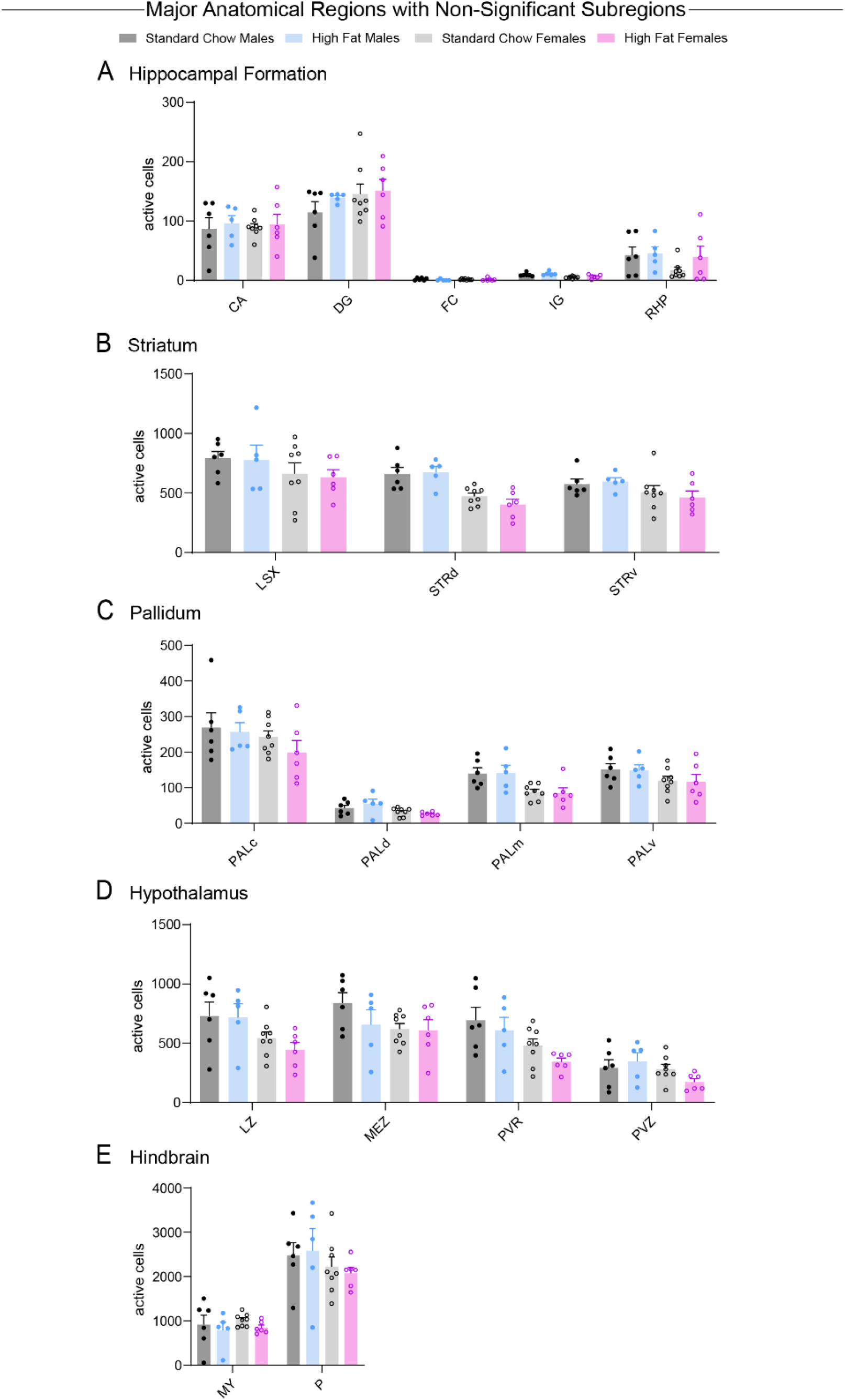
Active population cell quantification within major anatomical regions that do not contain significant subregions. A Hippocampal Formation (F(12, 105) = 0.7873, p = 0.6624). B Striatum (F(6, 63) = 0.3748, p = 0.8923). C Pallidum (F(9, 84) = 0.4681, p = 0.8921). D Hypothalamus (F(9, 84) = 0.6507, p = 0.7505). E Hindbrain (F(3, 42) = 0.7684, p = 0.5182). Data are shown as mean ± s.e.m. *p < 0.05, **p < 0.01**, *p < 0.001, ****p < 0.0001, two-way ANOVA with Šídák’s multiple comparison post hoc test.

**Supplemental Figure 4.**
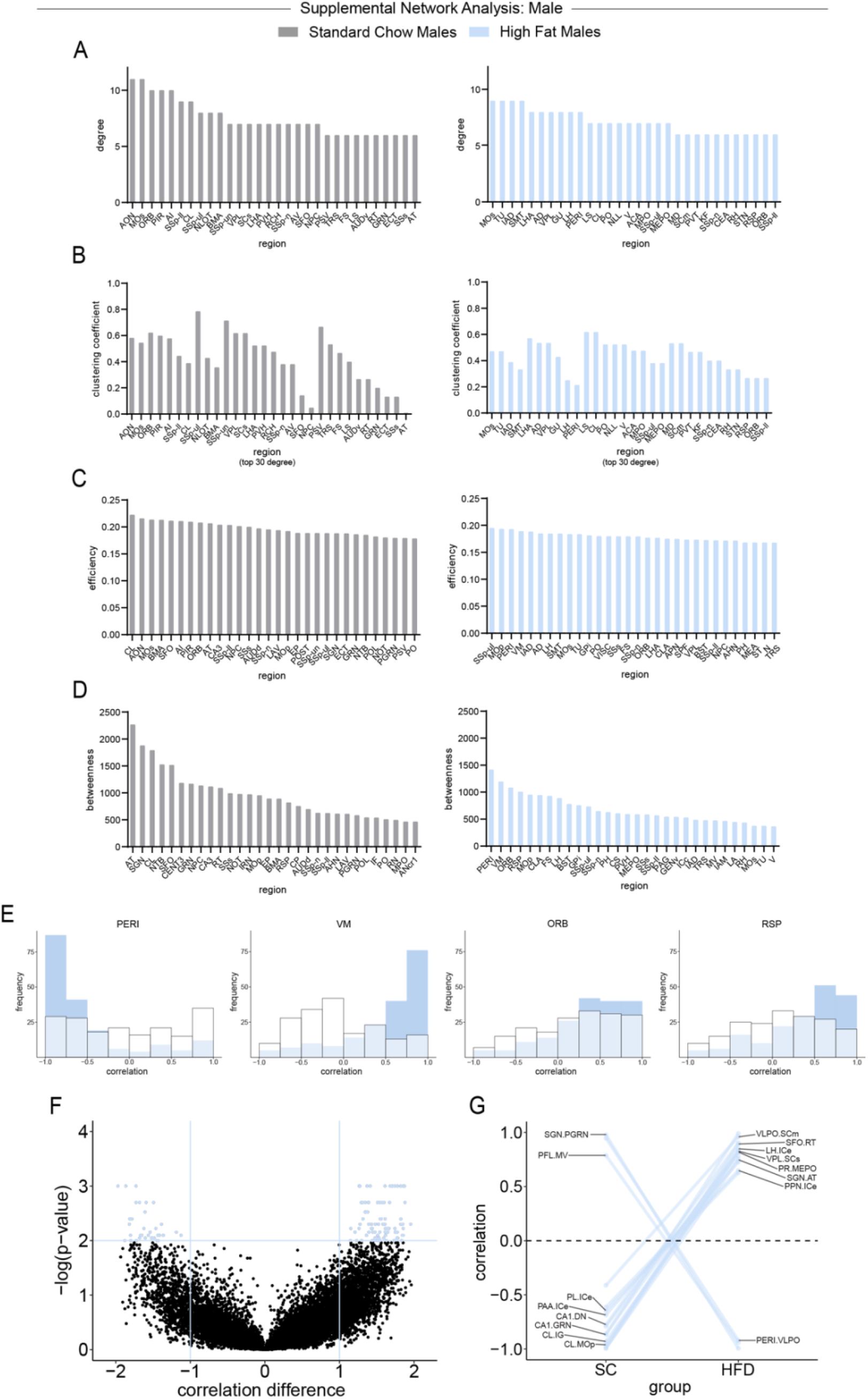
Male whole-brain correlational analysis of standard chow (gray) and high fat diet (blue) by utilizing SMARTTR. A Top 30 degree regions. B Clustering coefficient of top 30 degree regions. C Top 30 efficiency regions. D Top 30 betweenness regions. E Standard chow (white) and high fat diet (blue) Pearson-r correlation distributions of top 4 high fat diet betweenness regions. F Permutation difference analysis of each standard chow correlation to its identical high fat diet correlation pairing (sig. α = 0.01, blue; negative = 34, positive = 84). G Visualization of largest correlational changes in permutation analysis between standard chow and high fat diet (α = 0.001).

**Supplemental Figure 5.**
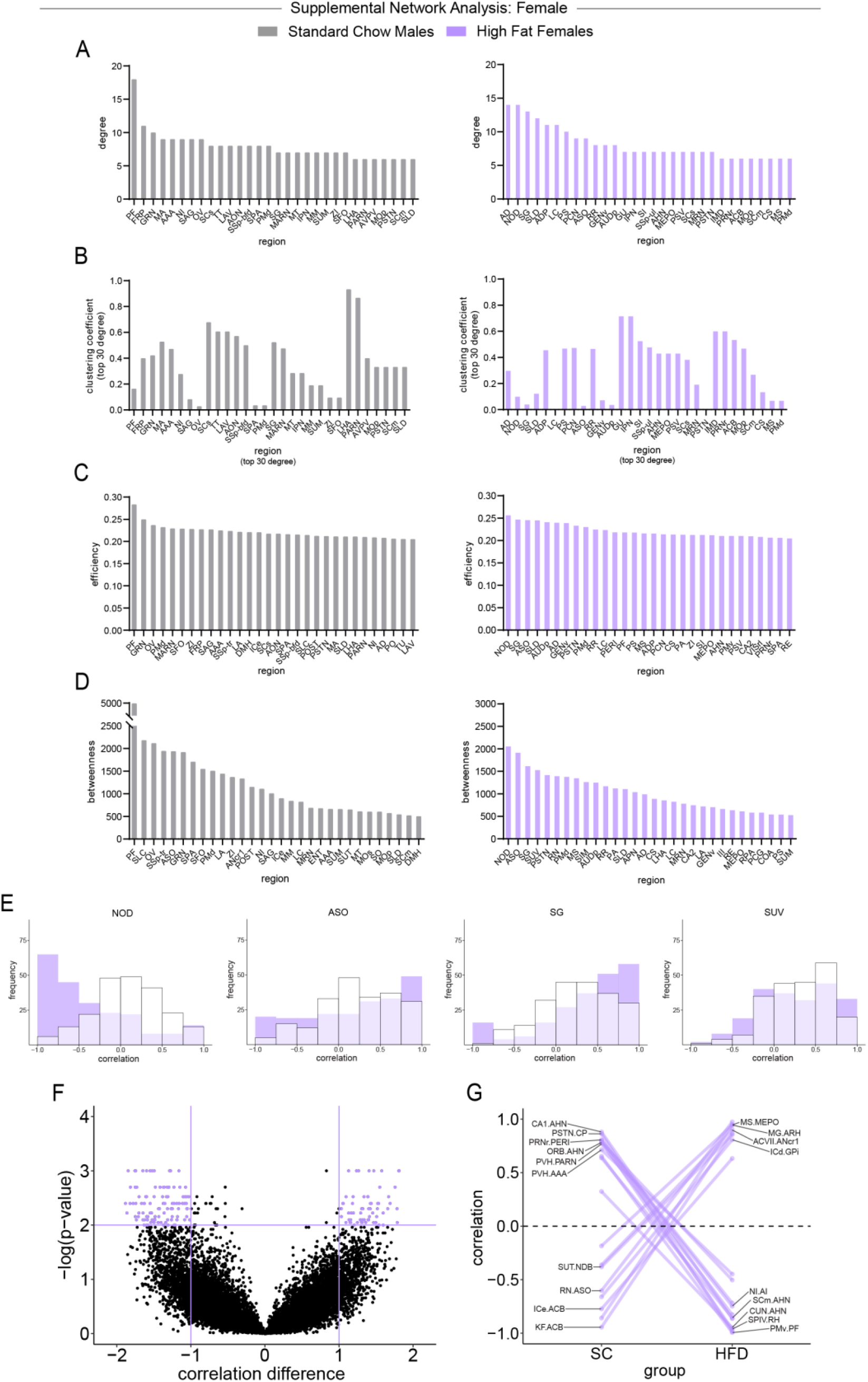
Female whole-brain correlational analysis of standard chow (gray) and high fat diet (purple) by utilizing SMARTTR. A Top 30 degree regions. B Clustering coefficient of top 30 degree regions. C Top 30 efficiency regions. D Top 30 betweenness regions. E Standard chow (white) and high fat diet (purple) Pearson-r correlation distributions of top 4 high fat diet betweenness regions. F Permutation difference analysis of each standard chow correlation to its identical high fat diet correlation pairing (sig. α = 0.01, purple; negative = 128, positive = 66). G Visualization of largest correlational changes in permutation analysis between standard chow and high fat diet (α = 0.001).

