## Supplemental Table and Figures for "Brain-wide Activity Mapping Reveals the Somatosensory Cortex as a Sex-Specific Regulator of High-Fat Diet Intake"

### SUPPLEMENTAL MATERIALS

**Table 1.** Allen Brain Atlas Common Coordinate Framework regions of interest list with acronym and full name.

***Table 1** Allen Brain Atlas Common Coordinate Framework*

| | | | |
|---|---|---|---|
| AAA - Anterior amygdalar area | GPi - Globus pallidus, internal segment | NOD - Nodulus (X) | SAG - Nucleus sagulum |
| ACA - Anterior cingulate area | GRN - Gigantocellular reticular nucleus | NOT - Nucleus of the optic tract | sAMY - Striatum-like amygdalar nuclei |
| ACB - Nucleus accumbens | GU - Gustatory areas | NPC - Nucleus of the posterior commissure | SBPV - Subparaventricular zone |
| ACVII - Accessory facial motor nucleus | IA - Intercalated amygdalar nucleus | NTB - Nucleus of the trapezoid body | SCH - Suprachiasmatic nucleus |
| AD - Anterodorsal nucleus | IAD - Interanterodorsal nucleus of the thalamus | NTS - Nucleus of the solitary tract | SCO - Subcommissural organ |
| ADP - Anterodorsal preoptic nucleus | IAM - Interanteromedial nucleus of the thalamus | ORB - Orbital area | SCm - Superior colliculus, motor related |
| AHN - Anterior hypothalamic nucleus | IC - Inferior colliculus | OT - Olfactory tubercle | SCs - Superior colliculus, sensory related |
| AI - Agranular insular area | ICB - Infracerebellar nucleus | OV - Vascular organ of the lamina terminalis | SF - Septofimbrial nucleus |
| AM - Anteromedial nucleus | ICc - Inferior colliculus, central nucleus | P - Pons | SFO - Subfornical organ |
| ANcr1 - Crus 1 | ICd - Inferior colliculus, dorsal nucleus | PA - Posterior amygdalar nucleus | SG - Supragenual nucleus |
| ANcr2 - Crus 2 | ICe - Inferior colliculus, external nucleus | PAA - Piriform-amygdalar area | SGN - Suprageniculate nucleus |
| AOB - Accessory olfactory bulb | IF - Interfascicular nucleus raphe | PAG - Periaqueductal gray | SI - Substantia innominata |
| AON - Anterior olfactory nucleus | IG - Induseum griseum | PALc - Pallidum, caudal region | SIM - Simple lobule |
| APN - Anterior pretectal nucleus | III - Oculomotor nucleus | PALd - Pallidum, dorsal region | SLC - Subceruleus nucleus |
| ARH - Arcuate hypothalamic nucleus | ILA - Infralimbic area | PALm - Pallidum, medial region | SLD - Sublaterodorsal nucleus |
| ASO - Accessory supraoptic group | IMD - Intermediodorsal nucleus of the thalamus | PALv - Pallidum, ventral region | SMT - Submedial nucleus of the thalamus |
| AT - Anterior tegmental nucleus | IP - Interposed nucleus | PARN - Parvicellular reticular nucleus | SNc - Substantia nigra, compact part |
| AUD - Auditory areas | IPN - Interpeduncular nucleus | PBG - Parabigeminal nucleus | SNr - Substantia nigra, reticular part |
| AUDd - Dorsal auditory area | IRN - Intermediate reticular nucleus | PCG - Pontine central gray | SO - Supraoptic nucleus |
| AUDp - Primary auditory area | IV - Trochlear nucleus | PCN - Paracentral nucleus | SPA - Subparafascicular area |
| AUDv - Ventral auditory area | KF - Koelliker-Fuse subnucleus | PERI - Perirhinal area | SPF - Subparafascicular nucleus |
| AV - Anteroventral nucleus of thalamus | LA - Lateral amygdalar nucleus | PF - Parafascicular nucleus | SPIV - Spinal vestibular nucleus |
| AVP - Anteroventral preoptic nucleus | LAV - Lateral vestibular nucleus | PFL - Paraflocculus | SS - Somatosensory areas |
| AVPV - Anteroventral periventricular nucleus | LC - Locus ceruleus | PG - Pontine gray | SSp-bfd - Primary somatosensory area, barrel field |
| B - Barrington's nucleus | LD - Lateral dorsal nucleus of thalamus | PGRN - Paragigantocellular reticular nucleus | SSp-ll - Primary somatosensory area, lower limb |
| BLA - Basolateral amygdalar nucleus | LDT - Laterodorsal tegmental nucleus | PH - Posterior hypothalamic nucleus | SSp-m - Primary somatosensory area, mouth |
| BMA - Basomedial amygdalar nucleus | LH - Lateral habenula | PIR - Piriform area | SSp-n - Primary somatosensory area, nose |
| BST - Bed nuclei of the stria terminalis | LHA - Lateral hypothalamic area | PL - Prelimbic area | SSp-tr - Primary somatosensory area, trunk |
| CA - Ammon's horn | LING - Lingula (I) | PMd - Dorsal premammillary nucleus | SSp-ul - Primary somatosensory area, upper limb |
| CA1 - Field CA1 | LP - Lateral posterior nucleus of the thalamus | PMv - Ventral premammillary nucleus | SSp-un - Primary somatosensory area, unassigned |
| CA2 - Field CA2 | LPO - Lateral preoptic area | PN - Paranigral nucleus | SSs - Supplemental somatosensory area |
| CA3 - Field CA3 | LS - Lateral septal nucleus | PO - Posterior complex of the thalamus | STN - Subthalamic nucleus |
| CEA - Central amygdalar nucleus | LSX - Lateral septal complex | POL - Posterior limiting nucleus of the thalamus | STRd - Striatum dorsal region |
| CENT2 - Lobule II | LT - Lateral terminal nucleus of the accessory optic tract | POR - Superior olivary complex, periolivary region | STRv - Striatum ventral region |
| CENT3 - Lobule III | LZ - Hypothalamic lateral zone | POST - Postsubiculum | SUM - Supramammillary nucleus |
| CL - Central lateral nucleus of the thalamus | MA - Magnocellular nucleus | PP - Peripeduncular nucleus | SUT - Supratrigeminal nucleus |
| CLA - Claustrum | MA3 - Medial accesory oculomotor nucleus | PPN - Pedunculopontine nucleus | SUV - Superior vestibular nucleus |
| CLI - Central linear nucleus raphe | MARN - Magnocellular reticular nucleus | PPT - Posterior pretectal nucleus | TEa - Temporal association areas |
| CM - Central medial nucleus of the thalamus | MBmot - Midbrain, motor related | PR - Perireunensis nucleus | TM - Tuberomammillary nucleus |
| COA - Cortical amygdalar area | MBsen - Midbrain, sensory related | PRM - Paramedian lobule | TR - Postpiriform transition area |
| COPY - Copula pyramidis | MBsta - Midbrain, behavioral state related | PRNr - Pontine reticular nucleus | TRN - Tegmental reticular nucleus |
| CP - Caudoputamen | MD - Mediodorsal nucleus of thalamus | PRP - Nucleus prepositus | TRS - Triangular nucleus of septum |
| CS - Superior central nucleus raphe | MEA - Medial amygdalar nucleus | PRT - Pretectal region | TT - Taenia tecta |
| CUL4, 5 - Lobules IV-V | MEPO - Median preoptic nucleus | PS - Parastrial nucleus | TU - Tuberal nucleus |
| CUN - Cuneiform nucleus | MEV - Midbrain trigeminal nucleus | PSTN - Parasubthalamic nucleus | V - Motor nucleus of trigeminal |
| DCO - Dorsal cochlear nucleus | MEZ - Hypothalamic medial zone | PSV - Principal sensory nucleus of the trigeminal | VAL - Ventral anterior-lateral complex of the thalamus |
| DG - Dentate gyrus | MG - Medial geniculate complex | PT - Parataenial nucleus | VCO - Ventral cochlear nucleus |
| DMH - Dorsomedial nucleus of the hypothalamus | MH - Medial habenula | PVH - Paraventricular hypothalamic nucleus | VII - Facial motor nucleus |
| DN - Dentate nucleus | MM - Medial mammillary nucleus | PVR - Periventricular region | VIS - Visual areas |
| DORpm - Thalamus, polymodal association cortex related | MO - Somatomotor areas | PVT - Paraventricular nucleus of the thalamus | VISC - Visceral area |
| DORsm - Thalamus, sensory-motor cortex related | MOB - Main olfactory bulb | PVZ - Periventricular zone | VISa - Anterior area |
| DP - Dorsal peduncular area | MOp - Primary motor area | Pa4 - Paratrochlear nucleus | VISam - Anteromedial visual area |
| DR - Dorsal nucleus raphe | MOs - Secondary motor area | RAmb - Midbrain raphe nuclei | VISp - Primary visual area |
| DT - Dorsal terminal nucleus of the accessory optic tract | MPN - Medial preoptic nucleus | RCH - Retrochiasmatic area | VISpm - posteromedial visual area |
| DTN - Dorsal tegmental nucleus | MPO - Medial preoptic area | RE - Nucleus of reuniens | VISrl - Rostrolateral visual area |
| ECT - Ectorhinal area | MRN - Midbrain reticular nucleus | RH - Rhomboid nucleus | VLPO - Ventrolateral preoptic nucleus |
| ENT - Entorhinal area | MS - Medial septal nucleus | RHP - Retrohippocampal region | VM - Ventral medial nucleus of the thalamus |
| EP - Endopiriform nucleus | MT - Medial terminal nucleus of the accessory optic tract | RL - Rostral linear nucleus raphe | VMH - Ventromedial hypothalamic nucleus |
| EW - Edinger-Westphal nucleus | MV - Medial vestibular nucleus | RM - Nucleus raphe magnus | VPL - Ventral posterolateral nucleus of the thalamus |
| FL - Flocculus | MY - Medulla | RN - Red nucleus | VPM - Ventral posteromedial nucleus of the thalamus |
| FN - Fastigial nucleus | NB - Nucleus of the brachium of the inferior colliculus | RPA - Nucleus raphe pallidus | VTA - Ventral tegmental area |
| FRP - Frontal pole, cerebral cortex | NDB - Diagonal band nucleus | RPO - Nucleus raphe pontis | VTN - Ventral tegmental nucleus |
| FS - Fundus of striatum | NI - Nucleus incertus | RR - Midbrain reticular nucleus, retrorubral area | ZI - Zona incerta |
| GENv - Geniculate group, ventral thalamus | NLL - Nucleus of the lateral lemniscus | RSP - Retrosplenial area | |
| GPe - Globus pallidus, external segment | NLOT - Nucleus of the lateral olfactory tract | RT - Reticular nucleus of the thalamus | |

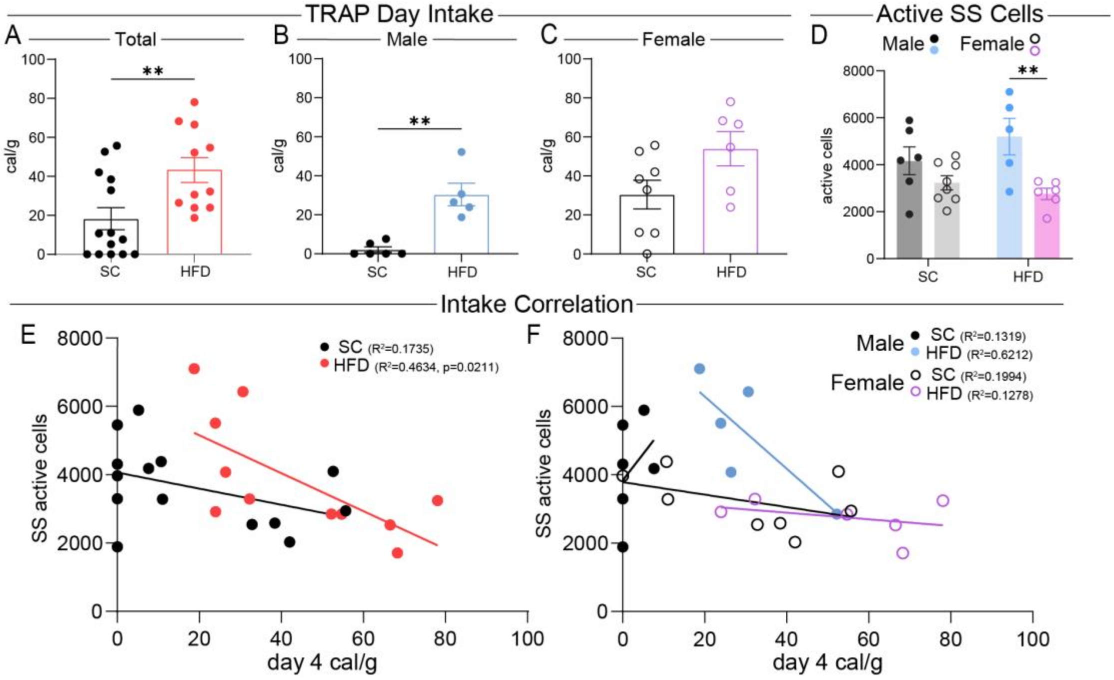


**Supplemental Figure 1.** Targeted recombination in active populations consumption and activity correlations. Standard chow (black) and high fat diet (colored) day 4 consumption in cal/g for A total (t = 2.921, df = 23, p = 0.0077), B male (t = 5.186, df = 9, p = 0.0006), and C female (t = 2.066, df = 12, p = 0.0611). D Sex-delineated active somatosensory (SS) cells during day 4 consumption (F(3, 21) = 6.675, p = 0.0024). Intake correlation with active SS cells for E total (SC: $R^2 = 0.1735$, p = 0.1384; HFD: $R^2 = 0.4634$, p = 0.0211), F male (SC: $R^2 = 0.1319$, p = 0.4791; HFD: $R^2 = 0.6212$, p = 0.1133), and G female (SC: $R^2 = 0.1994$, p = 0.2674; HFD: $R^2 = 0.1278$, p = 0.4867). Data are shown as mean ± s.e.m. *p < 0.05, **p < 0.01,** *p < 0.001, ****p < 0.0001, one-way ANOVA, unpaired Student's t-test, and linear regression.

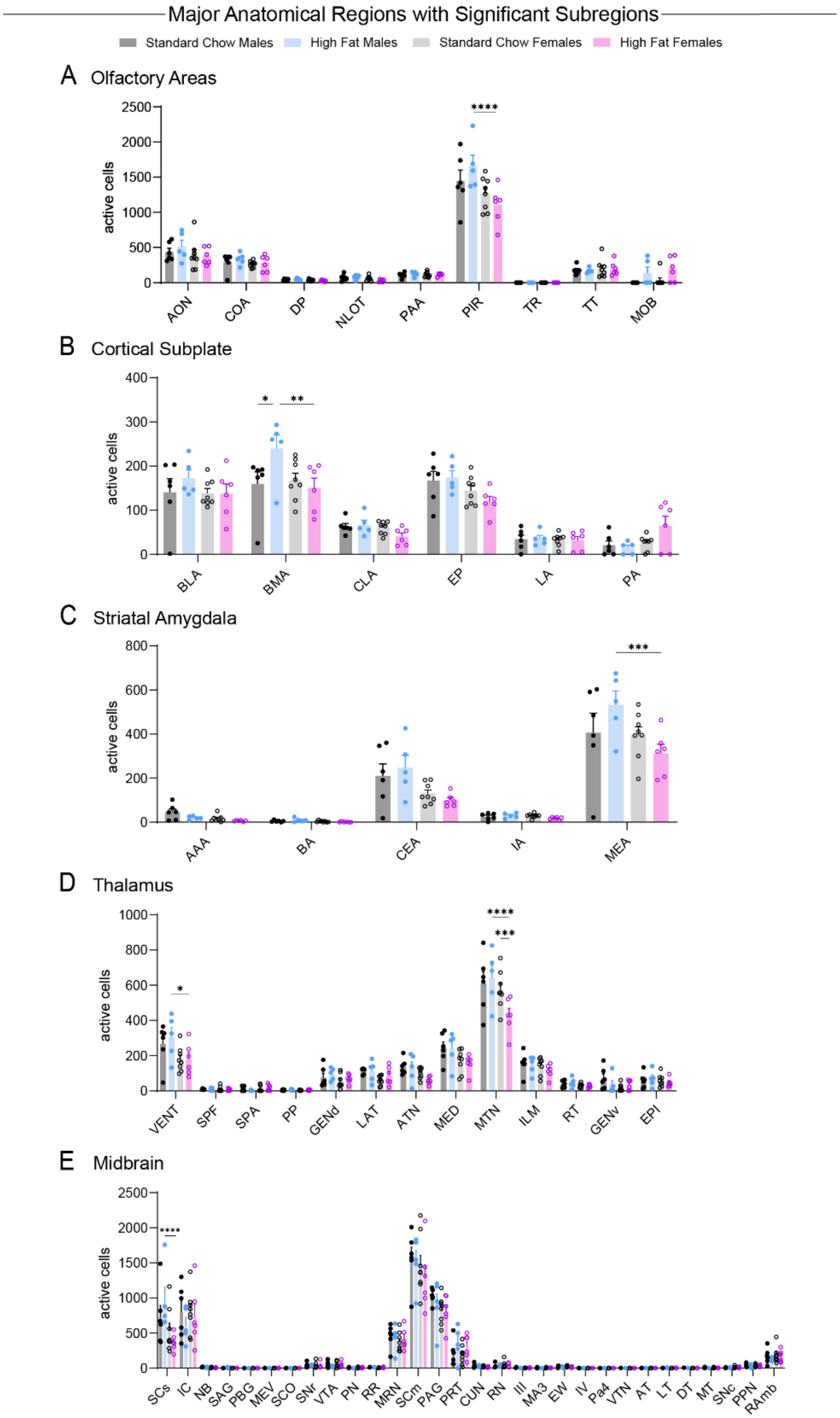


**Supplemental Figure 2.** Active population cell quantification within major anatomical regions that contain significant subregions. A Olfactory Areas ($F(24, 189) = 2.241$, $p = 0.0014$). B Cortical Subplate ($F(15, 126) = 1.736$, $p = 0.0519$). C Striatal Amygdala ($F(12, 105) = 1.723$, $p = 0.0720$). D Thalamus ($F(36, 273) = 1.698$, $p = 0.0102$). E Midbrain ($F(87, 630) = 1.069$, $p = 0.3238$). Data are shown as mean ± s.e.m. *$p < 0.05$, **$p < 0.01$,** *$p < 0.001$, ****$p < 0.0001$, two-way ANOVA with Šídák's multiple comparison post hoc test.

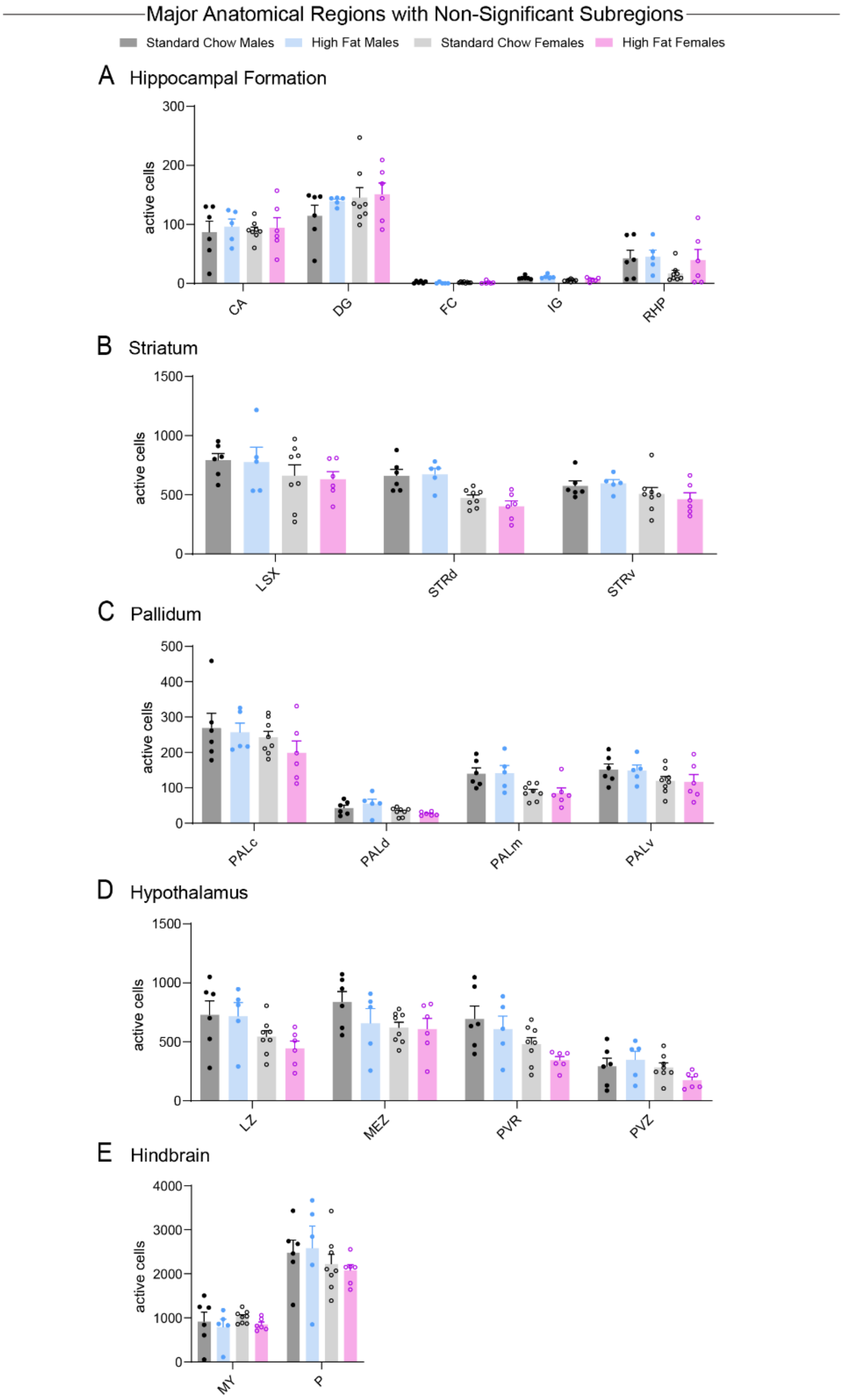


**Supplemental Figure 3.** Active population cell quantification within major anatomical regions that do not contain significant subregions. A Hippocampal Formation ($F(12, 105) = 0.7873$, $p = 0.6624$). B Striatum ($F(6, 63) = 0.3748$, $p = 0.8923$). C Pallidum ($F(9, 84) = 0.4681$, $p = 0.8921$). D Hypothalamus ($F(9, 84) = 0.6507$, $p = 0.7505$). E Hindbrain ($F(3, 42) = 0.7684$, $p = 0.5182$). Data are shown as mean ± s.e.m. *$p < 0.05$, **$p < 0.01$,** *$p < 0.001$, ****$p < 0.0001$, two-way ANOVA with Šídák's multiple comparison post hoc test.

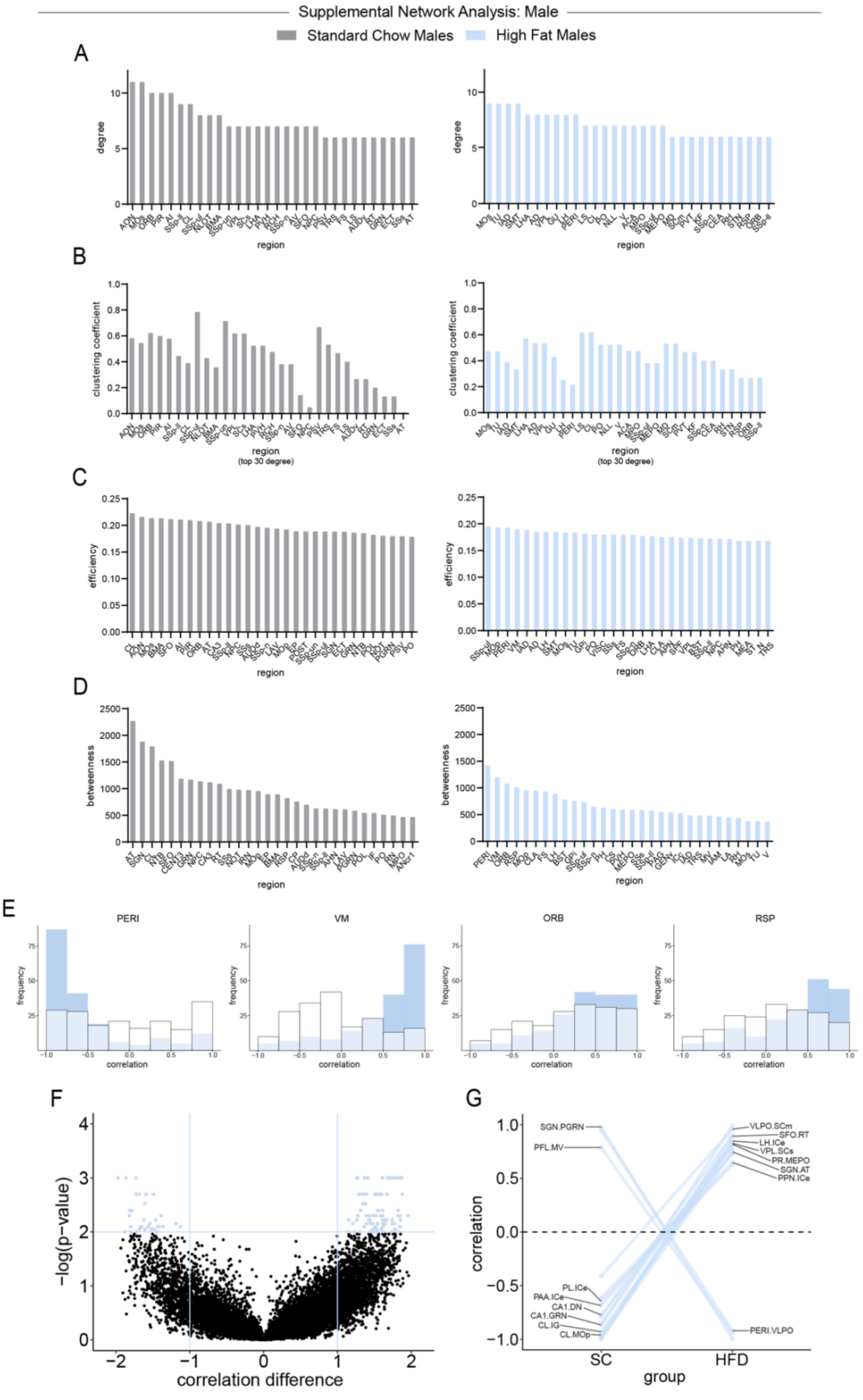


**Supplemental Figure 4.** Male whole-brain correlational analysis of standard chow (gray) and high fat diet (blue) by utilizing SMARTTR. A Top 30 degree regions. B Clustering coefficient of top 30 degree regions. C Top 30 efficiency regions. D Top 30 betweenness regions. E Standard chow (white) and high fat diet (blue) Pearson-r correlation distributions of top 4 high fat diet betweenness regions. F Permutation difference analysis of each standard chow correlation to its identical high fat diet correlation pairing (sig. $\alpha = 0.01$, blue; negative = 34, positive = 84). G Visualization of largest correlational changes in permutation analysis between standard chow and high fat diet ($\alpha = 0.001$).

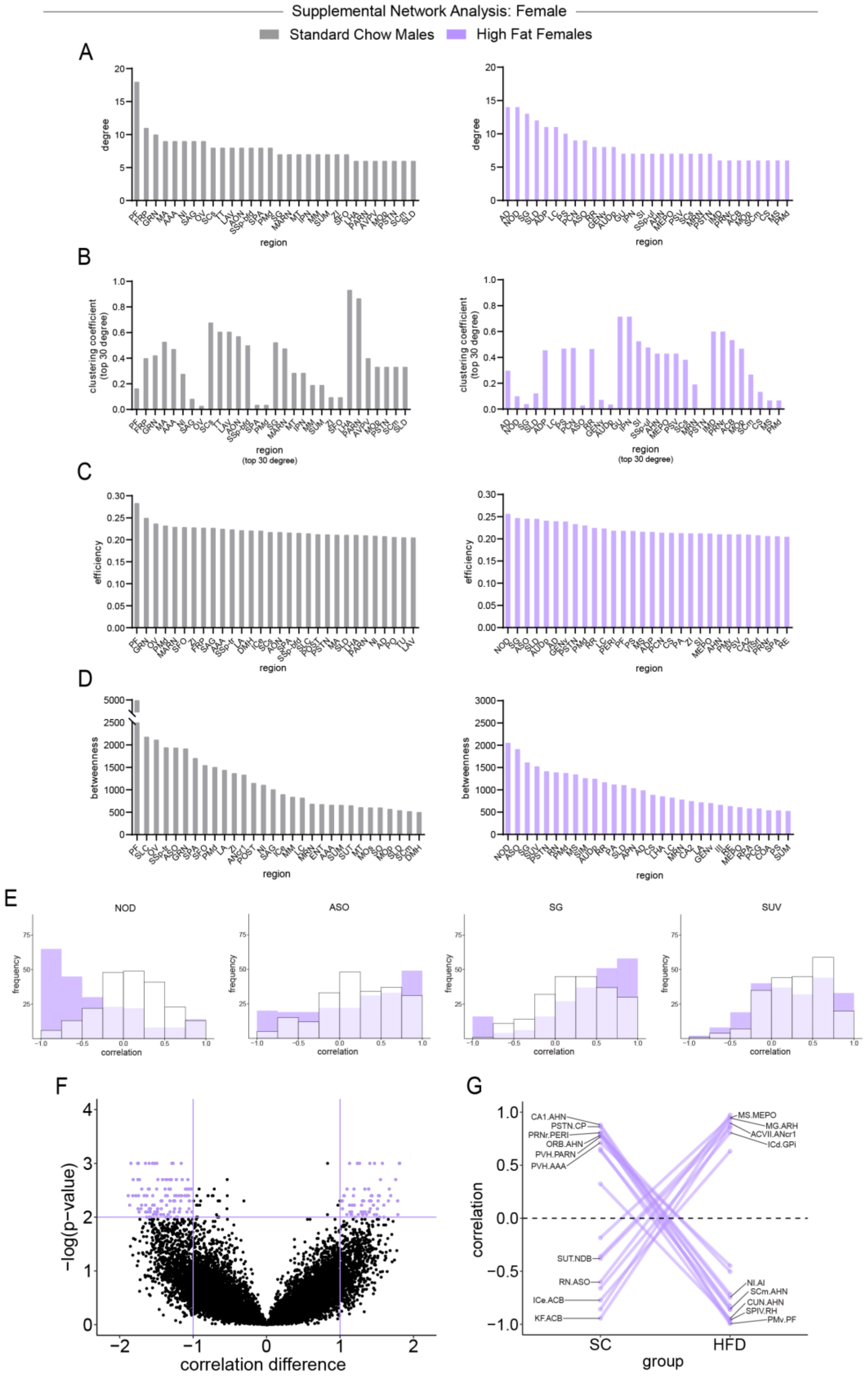


**Supplemental Figure 5.** Female whole-brain correlational analysis of standard chow (gray) and high fat diet (purple) by utilizing SMARTTR. A Top 30 degree regions. B Clustering coefficient of top 30 degree regions. C Top 30 efficiency regions. D Top 30 betweenness regions. E Standard chow (white) and high fat diet (purple) Pearson-r correlation distributions of top 4 high fat diet betweenness regions. F Permutation difference analysis of each standard chow correlation to its identical high fat diet correlation pairing (sig. α = 0.01, purple; negative = 128, positive = 66). G Visualization of largest correlational changes in permutation analysis between standard chow and high fat diet (α = 0.001).
